# Funneliformis mosseae and Pseudomonas putida-symbiotic interaction promote drought resilience in Citrus reticulata

**DOI:** 10.64898/2026.03.12.711468

**Authors:** Saleem Uddin, Sadia Gull, Jie Wang, Jinhan Yin, Hafiz Athar Hussain, Umer Mahmood, Xiaohong Yang

## Abstract

Climate change and increasing drought conditions significantly impedes citrus productivity in subtropical and tropical regions. This study explores the potential of combining arbuscular mycorrhizal fungi (AMF) *Funneliformis mosseae* and plant growth-promoting rhizobacteria (PGPR) *Pseudomonas putida* to mitigate drought resilience in *Citrus reticulata* (Red tangerine). AMF-mediated drought tolerance has been extensively documented; however, the collegial influence of PGPR and AMF on phytohormone signaling, photosynthetic efficiency, nutrient acquisition, and gene expression remains largely unexplored in citrus. We conducted a greenhouse experiment under both well water and drought stress conditions to assess the physiological and molecular responses to individual and co-inoculation with PGPR and AMF. Drought-stressed citrus plants, inoculated with AMF and PGPR, demonstrated significantly improved leaf water potential, stomatal conductance, carbon assimilation, and antioxidant defense. PGPR-AMF co-inoculation enhanced chlorophyll stability, osmotic adjustment, and nutrient uptake, while significantly reducing lipid peroxidation and ROS accumulation. The turquoise module emerged from transcriptomic and gene co-expression network analysis (WGCNA) as a potential key regulator of stress adaptation, revealed key regulatory transcription factors, e.g., *CrMYB4*, *CrZFP8*, *CrSOS5*, *CrRGFR2*, and *CrQUA1*, that were upregulated under combined inoculation, highlighting their potential role in stress adaptation. Our findings demonstrate that the synergistic PGPR-AMF interaction improves antioxidant enzyme activities and modulates gene expression to promote drought tolerance, providing new insights into the microbiome’s role in plant resilience. These results offer a potential strategy to boost citrus growth and yield under water scarcity, with broad implications for agricultural resilience to climate change.

## Introduction

As climate change exacerbates drought stress, posing significant risks to global productivity and environmental health (Zhou *et al*., 2019; Metze *et al*., 2023), mitigating these challenges has become a critical priority for food security (Mondal *et al*., 2023). However, plants do not function in isolation; they exist within intricate, co-evolved relationships with soil and plant-associated microorganisms that enhance growth, stress resilience, and adaptive responses to abiotic challenges (Pirozynski & Malloch, 1975; Hassani *et al*., 2018; Liu, H *et al*., 2024). These microbiomes provide essential nutrients, metabolites, and phytohormones, modulate plant defenses through functional barriers that mitigate environmental stress, offering promising strategies for combating drought (De Vries *et al*., 2020; Qi *et al*., 2022; Benmrid *et al*., 2023). Drought induces a range of metabolic changes, such as lower photosynthetic rates, modified root exudates, and increased reactive oxygen species (ROS), which in turn shift the microbiome in the rhizosphere through a phenomenon known as the “cry for help” hypothesis (Wang & Song, 2022; Lin *et al*., 2023; Metze *et al*., 2023). This response enriches beneficial microorganisms, such as *Streptomyces*, *Bacillus*, and *Enterococcus*, while decreasing microbial diversity (Francis *et al*., 2010; Santos-Medellín *et al*., 2021). The plant–microbiome dynamics under drought are further influenced by soil properties, particularly its water-holding capacity (De Vries *et al*., 2020; Allsup *et al*., 2023). Despite the emerging understanding of microbial roles, the underlying regulatory mechanisms of these shifts and their impact on plant health and productivity remain underexplored, highlighting the need for further investigation into these interactions for sustainable agricultural practices.

Citrus is a major fruit crop grown in over 140 countries, with China ranking as the largest global producer (Pan *et al*., 2019). In regions such as the sloped terrain of the Three Gorges Reservoir area, offers favorable conditions for high-quality citrus cultivation (Zeng *et al*., 2019). However, water scarcity poses a significant challenge, impacting root development, nutrient uptake, and photosynthesis, ultimately reducing growth and fruit quality (Rao *et al*., 2024). Drought, a major limiting factor for citrus production, adversely affects its physiology and productivity across tropical and subtropical regions (Chica & Albrigo, 2013; Wu *et al*., 2018; de Oliveira Sousa *et al*., 2022; Lu *et al*., 2023). With these challenges expected to worsen, developing sustainable strategies, such as utilizing beneficial microorganisms, is essential to enhance drought resistance in citrus and ensure long-term crop viability (Ngumbi & Kloepper, 2016).

Synthetic microbial communities (SynComs), combining AMF and PGPR, enhance drought tolerance in plants by improving water uptake, photosynthesis, nutrient acquisition, and oxidative stress responses, offering an eco-friendly substitute for chemical fertilizers and promoting resilience in crops like wheat, tomato, chickpea, and lettuce leaves (Gao *et al*., 2020; Marín *et al*., 2021; Saleem *et al*., 2021; Pradhan *et al*., 2022; Ikan *et al*., 2023). AMF symbioses facilitate plant access to water and nutrients by forming an intraradical mycelial network, which exchanges material and information between the plant and fungi (Lanfranco *et al*., 2016). This network regulates plant water uptake through modulation of aquaporin gene expression, facilitating water transport across air gaps under drought (Li *et al*., 2013). Additionally, AMF improve root system architecture (RSA) and support nodulation in legumes (Wang *et al*., 2021), and biomass in the tea plant (Liu, C-Y *et al*., 2024), thereby improving water absorption (Pan *et al*., 2020; Chandra *et al*., 2022). Furthermore, AMF enhance nutrient acquisition and drought tolerance by regulating the key signaling pathways such as *GmSPX5* in soybean, *SlSPX1* in tomato and the nitrate transporter gene *OsNPF4.5* in rice and increasing antioxidant activities (He *et al*., 2020; Wang *et al*., 2020; Liao *et al*., 2022; Yang *et al*., 2025). It also regulates plant hormonal balance, particularly by upregulating ABA and JA production, which together improve stress resilience (Duc *et al*., 2023; Ye *et al*., 2023). Through these mechanisms, AMF inoculation boosts drought resistance across diverse crops like maize, wheat, soybean, and citrus (Begum *et al*., 2019; Soliman *et al*., 2025; Zou *et al*., 2025).

Similarly, PGPR strains like *Paenibacillus polymyxa*, *Azospirillum brasilense*, *Pseudomonas sp*., and *Bacillus* improve drought tolerance by enhancing root architecture, nutrient uptake, and water retention in *Arabidopsis thaliana* (Yang *et al*., 2021), tomato (Gowtham *et al*., 2020), wheat (Barnawal *et al*., 2017), and maize (Yasmin *et al*., 2021; Li *et al*., 2025). Inoculation of *Pseudomonas putida* MTCC5279, *P. putida* NBRIRA and *B. amyloliquefaciens* NBRISN13 has been shown to increase root length, shoot growth and productivity, lateral root density, and nodule formation, thereby promoting nitrogen fixation and improving plant biomass under water-limited conditions in chickpea (Kumar *et al*., 2016; Tiwari *et al*., 2016). Additionally, PGPR modulate phytohormone synthesis, regulate stress-responsive gene expression to mitigate oxidative stress and enhance drought adaptation (Vejan *et al*., 2016; Liu, F *et al*., 2023). *Pseudomonas putida* FBKV2 enhances drought tolerance by increasing relative water content, proline, and chlorophyll content, while reducing MDA accumulation (Sandhya *et al*., 2009). Additionally, it enhances osmotic and water potential, chlorophyll, osmolyte content, and antioxidant enzyme activity, improving drought resistance (Ilyas *et al*., 2020; He *et al*., 2021). Overall, PGPR is an effective biofertilizer to enhance plant growth, nutrient uptake, and drought resilience, presenting an eco-friendly fertilizers (Azeem *et al*., 2022; Kour & Yadav, 2022).

Building upon the understanding that drought induces complex physiological and molecular changes in plants, this study hypothesizes that inoculation of *Funneliformis mosseae* and *Pseudomonas putida* enhances plant drought tolerance in red tangerine (*Citrus reticulata* Blanco) by modulating antioxidant activity, phytohormone regulation, nutrient acquisition mechanisms, and gene expression. These enriched microorganisms increase the plant resistance to water deficiency. The objectives of this study were to: (1) explore how these microbiota influence plant morphology, photosynthetic parameters, antioxidant activities, nutrient uptake, phytohormonal signaling, and gene expression under drought conditions; (2) By integrating advanced transcriptomic approaches, including WGCNA, this study aims to identify key stress-related genes modulated by AMF and PGPR interactions, thereby providing mechanistic insights into drought-resilient plant-microbe symbioses. This research uncovers potential novel perspectives on the contribution of combined microbial inoculation to drought resilience in citrus.

## Materials and Methods

### Plant material and growth conditions

Red Tangerine (*Citrus reticulata* Blanco), a widely used citrus rootstock, was used in this study. Seeds were provided by the Citrus Research Institute, Southwest University, Chongqing, China. To ensure surface sterility, seeds were treated with 70% ethanol for 30 seconds, followed by three rinses with distilled water and a treatment with a 0.5 M sodium hydroxide (NaOH) solution. Sterilized seeds were germinated in autoclaved river sand (particle size ≤ 2 mm) in a growth chamber at (28/20°C day/night; 80% relative humidity) to facilitate germination. After 4 weeks, uniform seedlings (∼4 cm height, four true leaves) were transplanted into 1.8 L plastic pots (top diameter 15.5 cm, bottom diameter 13.5 cm, height 11.5 cm) containing a sterilized soil (1.5 kg): sand mixture (v/v=5:2). Soil physicochemical properties were: total nitrogen (0.071 g kg^−1^), organic matter (14.87 g kg^−1^), available phosphorus (17.81 mg kg^−1^), available potassium (63.48 mg kg^−1^), pH (6.59), and alkali-hydrolyzable nitrogen (95.83 mg kg^−1^). Plants were supplied weekly with 10 mL of Hewitt’s nutrient solution lacking nitrogen and phosphorus (Hewitt, 1965). Plants were grown under greenhouse conditions with a photosynthetic photon flux density of 728-965 μmol/m^2^/s, an ambient temperature of 28.2°C/20.2°C (day/night), and a relative humidity of 75-85%. Pots were repositioned weekly to minimize positional bias and maintain uniform environmental conditions.

A completely randomized experimental design was employed with eight treatments; control (CK, no stress), control with AMF (+AMF), control with PGPR (+PGPR), control with both AMF and PGPR (AMF+PGPR), drought (D), drought with AMF (D+AMF), drought with PGPR (D+PGPR), and drought with AMF and PGPR (D+AMF+PGPR) (Fig. 1A). A total of three replicates per treatment were used, with ten plants assigned to each replicate (one seedling per pot), totaling 30 pots across all treatments.

**Fig. 1.**
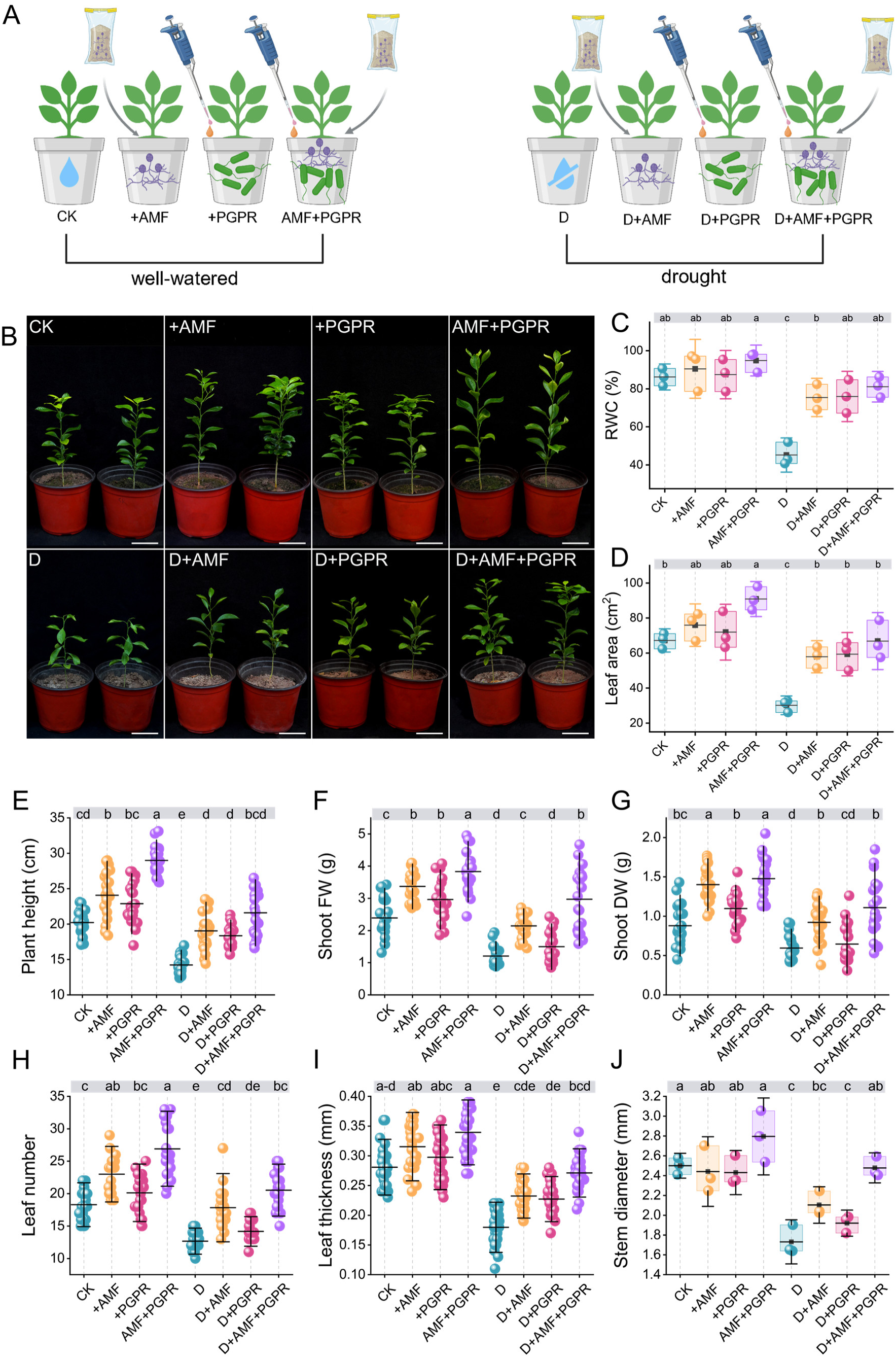
Effects of AMF and PGPR on shoot morphology and development under drought in red tangerine. **(A)** Experimental set-up. *Citrus reticulata* rootstock were inoculated with *F. mosseae* and *P. putida,* or without inoculation. **(B)** Representative images of *C. reticulata* ‘Red Tangerine’ (RT) after 70 days of treatment under well-watered (control, CK), AMF, PGPR, AMF+PGPR, drought stress (45% field capacity, D), D+AMF, D+PGPR, and D+AMF+PGPR conditions. **(C-J)** Morpho-physiological parameters of RT seedlings under the specified treatments: **(C)** Relative water content (RWC, %), **(D)** Leaf area (cm²), **(E)** Plant height (cm), **(F)** Shoot fresh weight (g), **(G)** Shoot dry weight (g), **(H)** Leaf number, **(I)** Leaf thickness (mm), and **(J)** Stem diameter (mm). Scale bar: 5 cm. Data represent means ± SD (*n* = 3 biological replicates, 25 plants per treatment per replicate). Means followed by different lowercase letters differ significantly among treatments (*P* < 0.05, Tukey’s HSD test following ANOVA).

### Microbial inoculation and drought stress application

The arbuscular mycorrhizal fungus strain *Funneliformis mosseae* (*F. mosseae*) was provided by Chongqing Engineering Research Center for Floriculture, Southwest University. The inoculum, consisting of root fragments, fungal spores, and hyphae, had a spore density of 12 spores/g and was propagated using maize (*Zea mays* L.). For both single and combined AMF + PGPR treatments, 50 g of inoculum was applied evenly to the seedling root zone in each pot.

The plant growth-promoting rhizobacterium *Pseudomonas putida* (GSICC 31637) was obtained from the Gansu Center of Industrial Culture Collection (GSICC), Lanzhou, China. The bacterial culture was maintained in Luria–Bertani (LB) medium at 30 °C for growth, with shaking until reaching 1 × 10^8 CFU mL^−1^. A 15 mL suspension was applied to the pots of each PGPR or AMF+PGPR treatment plant.

All plants were irrigated to 90% field capacity (FC) to allow acclimatization. After 20 days post-transplantation, on-stressed plants were maintained at 90% FC throughout the experiment, whereas drought-stressed plants were reduced to 45% FC by controlled water addition, with soil moisture monitored daily by pot weighing. Microbes were applied at the time of transplanting, and drought was imposed 20 days after transplanting. After 70 days of drought treatments, plants were harvested for growth, physiological, and transcriptomic analyses. Leaves were sampled from 15 plants in each treatment, with fully expanded leaves taken from the second and third intermediate nodes. Tissues from four plants were pooled per biological replicate, ground, and preserved at −80°C for subsequent analysis. Each treatment was evaluated using three biological replicates for transcriptomic profiling, physiological parameters, and growth traits.

### Plant growth, root system architecture, and physiological parameters

After 70 days of drought treatments, plants from each treatment were sampled to assess key growth parameters, including plant height, leaf thickness, leaf number, leaf area, shoot fresh and dry weight, stem diameter, and root fresh and dry weight. Root system architecture was analyzed using the Microtek ScanMaker i800 Plus imaging system, paired with WinRHIZO software (v2007b), which provided measurements of total root length, root surface area, mean root diameter, and root volume.

For Chlorophyll content, 100 mg of fresh leaf tissue was finely pulverized and homogenized in 10 mL of 80% (v/v) acetone (Chou *et al*., 2020). Overnight incubation of the homogenates was carried out at 25°C room temperature in the dark on a rotary shaker to ensure thorough pigment extraction. The samples were then centrifuged at 13,523 × g for 10 minutes, followed by the dilution of the resulting supernatants with an additional 10 mL of 80% acetone. Triplicate measurements of chlorophyll a and b concentrations were performed using a multimode microplate reader (Victor Nivo, PerkinElmer, UK) by recording absorbance at 663 nm, 645 nm, and 450 nm.

Relative water content (RWC) was determined by using the Barr and Weatherley method (Barrs & Weatherley, 1962). Fresh weight (FW) of the leaves was taken promptly after sampling. Samples were then rehydrated by immersing them in distilled water at 4 °C for 24 h in the dark to obtain the turgid weight (TW). The samples were over-dried at 70 °C to a constant weight to obtain dry weight (DW). RWC was computed as:

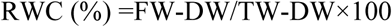

### Root mycorrhizal staining

Six plants from each treatment were randomly sampled. From each plant, approximately 14 root segments (∼1 cm in length) were excised for mycorrhizal staining. AMF colonization was evaluated using the trypan blue staining method described by Phillips and Hayman (Phillips & Hayman, 1970). Root segments were sequentially cleared in 10% KOH, bleached in 10% H₂O₂, acidified with 0.2 mol L⁻¹ HCl, and stained with 0.05% trypan blue. The degree of AMF colonization in roots was measured by using a ZEISS Axioskop 40 microscope (Carl Zeiss, Oberkochen, Germany).

### Determination of photosynthetic efficiency and stomatal morphology

Gas exchange parameters of citrus leaves were evaluated using a portable infrared gas analyzer (IRGA, LI-6400XT, LI-COR Biosciences, Lincoln, NE, USA). Fully expanded third and fourth leaves from each plant were sampled between 9:00 and 11:00 AM on clear days to assess net photosynthetic rate (Pn), transpiration rate (Tr), stomatal conductance (Gs), and intercellular CO₂ concentration (Ci).

For stomatal morphology, fully expanded second or third leaves from the apex were collected from each treatment. Small sections (∼5 × 5 mm) were excised from the central lamina, immediately immersed in 3% glutaraldehyde in 0.1 M phosphate buffer (pH 7.2), and fixed for 4 h at 4 °C. The samples were rinsed three times (10 min each) with 0.1 M phosphate buffer and dehydrated through an ethanol series (30%, 50%, 70%, 80%, and 100%), with two 10-min rinses each in absolute ethanol (100%). The dehydrated tissues were subsequently treated with isoamyl acetate substitution (50% once, 100% twice) and subjected to critical point drying using a CO₂ dryer (Eiko XD-1, Japan). Dried samples were affixed to aluminum stubs, coated with gold using an Eiko IB-3 (Japan) ion sputter coater, and examined with a JEOL JSM-840 scanning electron microscope (JEOL, Tokyo, Japan).

Stomatal density on the abaxial epidermis was calculated from Scanning Electron Microscopy (SEM) micrographs as stomata mm⁻². Stomatal pore width and length were measured using ImageJ (v. 1.53; NIH, USA), and aperture index was calculated as width/length ratio. Analyses covered ≥20 stomata treatment⁻¹ (*n* = 3).

### Reactive oxygen species (ROS) detection, Malondialdehyde (MDA) quantification, and antioxidant enzyme assays

Accumulation of hydrogen peroxide (H₂O₂) and superoxide anion (O₂⁻) in healthy and fully expanded *C. reticulata* leaves from each treatment were visualized histochemically via 3,3′-diaminobenzidine (DAB) and nitroblue tetrazolium (NBT) staining, respectively (*n* = 6 leaves /treatment⁻¹). Leaves were treated by vacuum infiltration with 10 mM potassium phosphate buffer (pH 7.8) for 30 min and subsequently incubated in 1 mg mL⁻¹ NBT or DAB at 25°C in darkness (NBT, 5 h; DAB, 12 h) as described by Khan et al (Khan *et al*., 2021). After staining, tissues were destained in boiling ethanol:acetic acid:glycerol (3:1:1, v/v/v) and photographed to assess ROS accumulation under AMF/PGPR and drought treatments.

The levels of antioxidant enzymes were observed with commercial assay kits (Nanjing Jiancheng Bioengineering Institute, Nanjing, China), as mentioned in the manufacturer’s guidelines. The enzymes and detection wavelengths were: MDA (A003-3-1) at 530 nm, superoxide dismutase (SOD) at 450 nm (A001-3), peroxidase (POD) at 420 nm (A084-3-1), glutathione reductase (GR) at 340 nm (A062-1-1), and catalase (CAT) at 405 nm (A007-1-1). The quantification of MDA content was analyzed using 0.1 g of homogenized leaf tissue extracted in 100 mM potassium phosphate buffer (pH 7.4), followed by centrifugation at 1205 g for 10 min at 4 °C to obtain the supernatant (Shen *et al*., 2021).

The activity of Ascorbate peroxidase (APX) determined spectrophotometrically based on the protocol mentioned by Yang *et al* (Yang *et al*., 2015). The assay mixture was composed of 50 mM phosphate buffer (pH 7.0), 2 mM ascorbate, 1 mM EDTA, and 0.1 mM H₂O₂. APX activity was measured by reduction in absorbance at 290 nm and expressed as units per milligram of protein (U mg⁻¹ protein), representing the enzymatic activity normalized to total protein content. All measurements were conducted using three independent biological replicates.

### Determination of plant nutrient concentrations

The concentrations of macronutrients (N, P, and K) and micronutrients (Fe, Mn, and Zn) in citrus leaves and roots were quantified after acid digestion and subsequent elemental analysis. For total nitrogen (N) determination, 0.2 g of oven-dried and finely ground leaf and root tissue was digested using a H₂SO₄-H₂O₂ mixture following the Kjeldahl method (Ogg, 1960). Organic nitrogen was converted to ammonium sulfate through digestion, and ammonia was subsequently distilled and titrated using a Kjeldahl nitrogen analyzer (KN-520, Alva, China). Nitrogen content was expressed on a dry-weight basis (g kg⁻¹ DW).

For P, K, Fe, Mn, and Zn analysis, 0.1 g of dried leaf and root powder was digested with concentrated HNO₃ into PTFE vessels, followed by stepwise heating at 80°C for 2 h, 120°C for 2 h, and 160°C for 4 h until complete digestion. The digests were evaporated gently to near dryness, diluted to 25 mL with 1% HNO₃, and filtered. Reagent blanks were prepared in parallel.

Elemental concentrations were quantified by inductively coupled plasma optical emission spectrometry (ICP-OES; iCAP 7200 HS Duo, Thermo Fisher Scientific, USA) and inductively coupled plasma mass spectrometry (ICP-MS; iCAP RQ, Thermo Fisher Scientific, USA), depending on the detection requirements. Quantification was based on external calibration using certified multi-element standards. Analytical precision and accuracy were ensured by running blanks and reference samples.

### Quantification of endogenous plant hormones

Endogenous phytohormones such as Indole-3-acetic Acid (IAA), Abscisic Acid (ABA), Jasmonic Acid (JA), 1-Aminocyclopropane-1-carboxylic Acid (ACC), *trans*-Zeatin Riboside (*t*ZR), and Gibberellin (GA₁) were quantified from each treatment using ultra-high-performance liquid chromatography–tandem mass spectrometry (UHPLC–MS/MS).

Leaf tissue (≈100 mg) was homogenized in liquid nitrogen, followed by extraction in 1 mL of isopropanol: water (v/v=80:20) containing isotopically labeled internal standards. After vortexing, homogenization, and ultrasonic treatment in an ice bath, the samples were incubated at −20 °C for 1 h and centrifuged at 14,000 × g for 15 min at 4 °C. The supernatant was dried to completion and reconstituted in methanol: water (v/v=50:50) before being filtered through a 0.22 μm PTFE membrane. Chromatographic separation was carried out on an HSS T3 column (100 × 2.1 mm, 1.8 μm) at 40°C, with a mobile phase of water (0.1% formic acid) and acetonitrile (0.1% formic acid) under a linear gradient. Multiple reaction monitoring (MRM) with electrospray ionization (ESI) in positive and negative modes was used for detection, while instrument control as well as data analysis were accomplished using Analyst v1.7.2 and Sciex OS v2.0.1.

Calibration curves were generated using mixed standards (0.01-50.00 ng/mL) with internal standard correction, and linearity was evaluated using the 1/x weighting method. Quantitative accuracy and precision were verified using spiked recovery and quality control samples, producing recoveries between 84% to 116% and RSDs <13%. The absolute concentrations, expressed in ng g^-^¹ fresh weight (FW), was determined using the following equation:

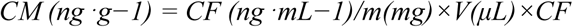

Where Cₘ is the metabolite concentration, C_F is the final measured concentration, V is the extraction volume, m is the sample mass, and CF is the concentration factor.

### RNA-Seq and bioinformatics analysis

RNA sequencing (RNA-seq) facilitated the identification of differentially expressed genes (DEGs) linked to drought stress. Citrus leaves exposed to drought, AMF, and PGPR treatments were used for total RNA extraction with TRIzol reagent (Invitrogen, USA). Each treatment was represented by three biological replicates, each consisting of three experimental repeats. Samples were assessed for RNA integrity using an Agilent 2100 Bioanalyzer (Agilent Technologies, USA), and those with RNA Integrity Number (RIN) ≥ 7.0 were selected for library construction. Sequencing libraries were prepared using the Illumina TruSeq RNA Sample Preparation Kit according to the manufacturer’s instructions and sequenced on an Illumina NovaSeq 6000 platform at Personalbio (Shanghai, China, project TR2024122010250ZH7) to generate 150 bp paired-end reads.

Raw reads were processed to remove adapter sequences and low-quality reads (Phred score < 20) using FASTP (Chen *et al*., 2018), yielding clean reads for downstream analysis. Over 99% of reads passed quality filtering, with GC contents ranging between 42% and 44% (Table S1). The high-quality reads were subsequently aligned to the *Citrus reticulata* (Wu *et al*., 2014) reference genome via HISAT2 (v2.2.1) (Kim *et al*., 2015) using default parameters. Transcript expression was quantified in terms of fragments per kilobase of transcript per million mapped reads (FPKM) with HTSeq. The biological replicates showed excellent reproducibility, as confirmed by correlation and principal component analyses (PCA) (Pearson’s r > 0.95). DEGs were identified using DESeq2 (v1.30.1) with a false discovery rate (FDR) < 0.05 and with fold change (FC) **≥** 1 as the selection criteria (Love *et al*., 2014). Heatmap visualization of the clustering results was carried out using R package heatmap (Kolde, 2019). Functional enrichment analyses of DEGs were performed using ClusterProfiler in R (v4.0), with Gene Ontology (GO) (http://www.geeontology.org) and Kyoto Encyclopedia of Genes and Genomes (KEGG) (http://www.kegg.jp), COG/KOG (https://www.ncbi.nlm.nih.gov/COG/), Pfam (http://pfam.xfam.org/), and Uniprot (http://www.uniprot.org/) databases as annotation sources. Enrichment was taken as significant at an adjusted *p*-value < 0.05 (Ashburner *et al*., 2000; Kanehisa & Goto, 2000).

### Weighted gene co-expression network analysis (WGCNA)

Transcriptomic data were analyzed to construct a weighted gene co-expression network using the Topological Overlap Matrix (TOM) similarity metric to quantify co-expression relationships among genes. To identify the gene regulatory network, the relationships between co-expression modules and physiological stress-responsive traits were examined with the WGCNA package (v1.72) in R (Langfelder & Horvath, 2008) utilizing normalized FPKM values higher than one. DEGs were further clustered using the R package Mfuzz (Kumar & Futschik, 2007) based on their normalized FPKM values, with three clusters defined to reveal distinct expression profiles among the treatments. A soft-thresholding power (β=15) was selected, as it corresponded to the point of stabilization in the scale-free topology fit, as depicted in the supplementary (Fig. S5). The soft threshold power was determined by pickSoftThreshold, which relied on a scale-free topology model fit (*R²*) greater than 0.9. The blockwiseModules function was applied to construct the co-expression network and to identify modules highly associated with the target trait. The analysis was performed using the following parameters: power = 18, minModuleSize = 50, TOM type = unsigned, maxBlockSize = 35,000, and mergeCutHeight = 0.25. Regulatory interactions between transcription factors and their co-expressed genes were predicted using the Plant Transcriptional Regulatory Map platform (PlantRegMap; http://plantregmap.gao-lab.org) (Tian *et al*., 2020). The resulting regulatory network was subsequently visualized in Cytoscape v3.9.0 (Smoot *et al*., 2011).

### Real-time qPCR validation

RT-qPCR analysis was performed on the remaining samples from RNA sequencing to validate the relative expression level of twelve hub genes via FastPure Universal Plant Total RNA Isolation Kit (Vazyme, China) as mentioned in the manufacturer’s protocol. Subsequently, reverse transcription of total RNA into cDNA was performed using the HiScript II Q RT SuperMix for qPCR Kit (R223-01; Vazyme, Nanjing, China). Quantitative real-time PCR was performed using the 2× Universal SYBR Green Fast qPCR Mix (RM21203; ABclonal, Wuhan, China). Each 10 μL reaction contained 1 μL of cDNA template, 5 μL of SYBR Green Master Mix, 0.2 μL of each primer (10 μM), and 3.6 μL of ultrapure water. The reaction procedure consisted of an initial denaturation at 95°C for 10 min, followed by 40 cycles of 95°C for 15s, 55-60°C for 15s, and 72°C for 15s, with a final extension at 72°C for 5 min. Annealing temperatures were optimized for individual primer pairs. Each sample included three biological and three technical replicates. The citrus β-actin gene was used as the internal control, and the target gene expression was quantified with the 2^⁻ΔΔCt^ method. Primer sequences are provided in Table S2.

## Statistical analysis

Statistical analysis were performed using Statistix 8.1 (Analytical Software, Tallahassee, FL, USA). Data were first tested for homogeneity of variance and, if necessary, transformed to meet ANOVA assumptions. Differences among treatment means were determined by one-way analysis of variance (ANOVA) followed by Tukey’s multiple comparison test at a significance level of p < 0.05. Mean values and their corresponding standard errors (± SE) were calculated from three independent replicates. Data visualization and graphical representations were generated using OriginPro 2024b (OriginLab, Northampton, MA, USA). Heatmaps and gene network visualizations were produced in R Studio (v2.4.3) and Cytoscape (v3.10.2), respectively.

## Results

### Growth and water status responses in citrus under drought and microbial inoculation

The influence of *F. mosseae* and *P. putida* on citrus drought tolerance was assessed 70 days post-inoculation by monitoring plant performance under both drought and control conditions. Phenotypic assessment (Fig. 1B) revealed pronounced differences in plant vigor, with drought-stressed plants exhibiting stunted growth and reduced foliage compared to controls. Inoculated plants, especially those with concurrent AMF+PGPR application, exhibited superior vigor and structural integrity under both regimes. Drought stress caused a significant decline in relative water content (RWC), leaf area, plant height, fresh and dry biomass, leaf number, leaf thickness, and stem diameter in rootstocks compared to the control.

Quantitative assessments revealed that drought stress markedly suppressed multiple growth traits in citrus seedlings. Relative water content (RWC) remained largely unchanged between the non-inoculated control (CK) and plants inoculated with AMF or PGPR alone under well-watered conditions. By contrast, combined AMF+PGPR inoculation significantly enhanced RWC by approximately 10% compared with CK. Under water deficit, RWC declined by 47.6% in non-inoculated plants, whereas inoculation substantially mitigated this reduction, with increases of 66%, 68%, and 79% recorded for AMF, PGPR, and AMF+PGPR treatments, respectively (Fig. 1C).

Leaf area was significantly enlarged in AMF, PGPR, and AMF+PGPR-treated plants under optimal irrigation compared with CK. Drought stress induced a sharp decline in leaf expansion, but inoculated seedlings maintained considerably greater leaf area than their non-inoculated counterparts (Fig. 1D). In parallel, drought treatment caused significant decreases in plant height (29.7%), shoot fresh weight (49.3%), shoot dry weight (32%), leaf number (30.7%), leaf thickness (35.7%), and stem diameter (30.8%) in CK seedlings (Fig. 1E-J). Inoculated plants under stress largely sustained biomass production, leaf number, tissue thickness, and stem diameter at levels comparable to non-stressed controls. Although height remained somewhat reduced in AMF and PGPR-treated plants under drought, dual inoculation (AMF+PGPR) not only prevented height loss but promoted significant increases of 43.4% under well-watered and 51.8% under drought conditions compared with CK. These findings demonstrate that microbial symbiosis, particularly AMF+PGPR co-inoculation, substantially enhances water conservation and growth resilience in citrus subjected to drought stress.

### Effects of microbial inoculation on root morphology under drought stress

Drought stress markedly suppressed root development in citrus rootstocks, as evidenced by reduced biomass and altered architecture (Fig. 2A). Root colonisation assays revealed that non-inoculated plants showed no root mycorrhizal colonisation. In contrast, colonisation with *F. mosseae* successfully established symbiosis under both water regimes, with the most extensive colonisation occurring in plants co-inoculated with AMF and PGPR (Fig. 2B, S1).

**Fig. 2.**
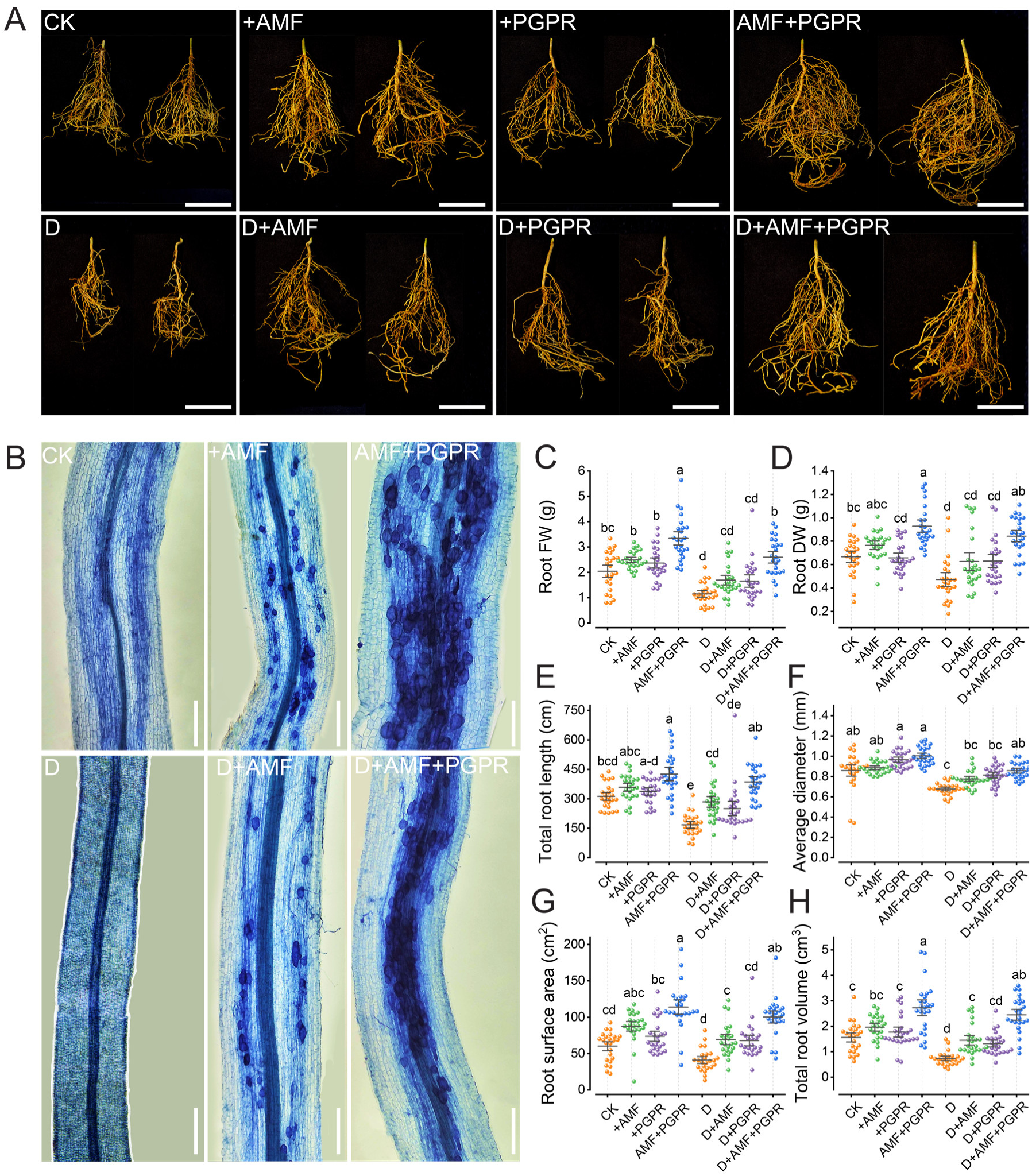
Effects of AMF and PGPR on root morphology and development under drought in red tangerine. **(A)** Representative images of whole root systems after 70 days under well-watered (control, CK), AMF, PGPR, AMF+PGPR, drought (45% field capacity, D), D+AMF, D+PGPR, and D+AMF+PGPR conditions. Scale bar = 5 cm. **(B)** Light microscopy images of AMF colonization patterns in RT roots under the respective treatments. Scale bar 100 μm. (C-H) Quantitative root morphological traits: **(C)** Root fresh weight (g), **(D)** Root dry weight (g), **(E)** Total root length (cm), **(F)** Average root diameter (mm), **(G)** Root surface area (cm²), **(H)** Total root volume (cm³). Scale bars: 5 cm and 100 μm. Data are means ± SD (*n* = 3 biological replicates, 25 plants per treatment per replicate). Different letters denote significant differences among treatments (*P* < 0.05, Tukey’s HSD test following ANOVA).

Quantitative assessments showed that water deficit reduced, non-inoculated plants exhibited pronounced reductions in root biomass, with fresh and dry weights declining by 43.9% and 29.8%, respectively, compared with the well-watered control. Solely inoculation with AMF and PGPR, and particularly with the AMF+PGPR combination, substantially mitigated these reductions; under well-watered conditions, this treatment enhanced root fresh weight to the highest level among all treatments, while under drought stress, it achieved 26.8% improvement relative to non-inoculated plants (Fig. 2C-D). Root architectural traits were also severely impaired by drought, with total root length, average diameter, surface area, and volume reduced by 46.6%, 20.9%, 31.5%, and 52.5%, respectively (Fig. 2E-H). In contrast, AMF+PGPR co-inoculation promoted robust root development, increasing total length, diameter, surface area, and volume by 36.1%, 16.2%, 89.5%, and 75% under well-watered conditions, and by 23.6%, 0.0%, 67%, and 57% under drought stress, respectively. The findings underscore how AMF and PGPR work synergistically to preserve root growth and architecture, thereby enhancing drought resilience in citrus.

### Stomatal pattern and leaf gas exchange in response to drought and microbial inoculation

Scanning electron microscopy (SEM) micrographics examination revealed distinct alterations in stomatal architecture across treatments, with drought-stressed non-inoculated plants (D) notably altered the appearance and exhibiting reduced stomatal density compared to controls in citrus leaves (Fig. 3A). Inoculation with *F. mosseae* and *P. putida*, either individually or combined (AMF+PGPR), restored stomatal density and morphology under both conditions, with the strongest effect observed in the AMF+PGPR co-inoculation (Fig. 3B). Stomatal aperture size also varied, with stressed controls showing narrower openings, while inoculated treatments, particularly AMF+PGPR co-inoculation, maintained wider apertures under drought (Fig. 3C).

**Fig. 3.**
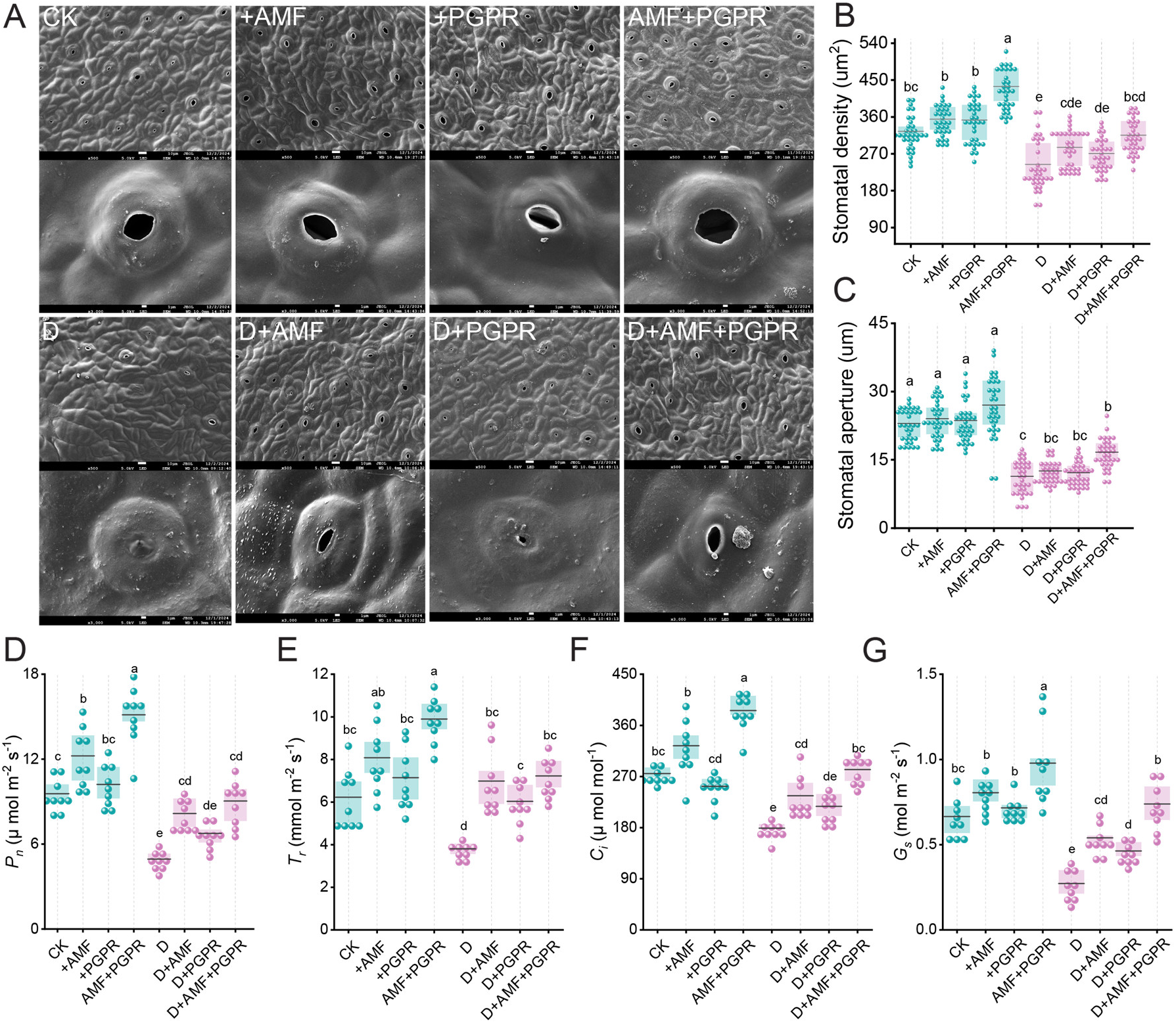
AMF and PGPR modulate stomatal morphology and gas exchange in red tangerine under drought stress. **(A)** Scanning electron micrographs of stomatal characteristics under well-watered (control, CK), AMF, PGPR, AMF+PGPR, drought, D+AMF, D+PGPR, and D+AMF+PGPR conditions. Top row: stomatal fields; bottom row: individual stomatal apertures. **(B-G)** Quantitative stomatal and physiological traits: **(B)** Stomatal density (stomata mm⁻²), **(C)** Stomatal aperture (µm), **(D)** Net photosynthetic rate (*Pn*, µmol m⁻² s⁻¹), **(E)** Transpiration rate (*Tr*, mmol m⁻² s⁻¹), **(F)** Intercellular CO₂ concentration (*Ci*, µmol mol⁻¹), **(G)** Stomatal conductance (*Gs*, mol m⁻² s⁻¹). Scale bar: 10 and 1 μm. Data are means ± SD (*n* = 3 biological replicates, 8–10 leaves per treatment per replicate). Significant differences among treatments are denoted by distinct letters (*P* < 0.05, Tukey’s HSD test following ANOVA).

Quantitative analysis confirmed that drought stress reduced stomatal density by approximately 24.5% in D relative to CK. Conversely, AMF, PGPR, and AMF+PGPR treatments under well water condition increased density by 9%, 8%, and 33.8%, respectively, with further enhancements of 12%, 17%, and 2% under water deficit (Fig. 3B). Stomatal aperture diminished by 50% in D, however, inoculation mitigated this reduction, with D+AMF+PGPR co-inoculation sustaining apertures 27% larger than D (Fig. 3C). Photosynthetic parameters were similarly affected. Net photosynthetic rate (*P_n_*) reduced by 48% in D, however AMF+PGPR co-inoculation nearly mitigate drought effects (Fig. 3D). Transpiration rate (*T_r_*) decreased by 39% in D, while inoculated plants showed increases of 12%, 3%, and 16% for AMF, PGPR, and AMF+PGPR co-inoculation, respectively, under drought (Fig. 3E). Internal CO₂ concentration (*C_i_*) dropped by 35% in D, but AMF+PGPR restored it to near-control levels with a 2.5% increase (Fig. 3F). Stomatal conductance (*G_s_*) was reduced by 59% in D, with D+AMF+PGPR not only recovered but enhanced *G_s_* by 11.15% compared to the control (Fig. 3G). These results underscore the pivotal role of microbial symbiosis, especially AMF+PGPR co-inoculation, in preserving stomatal function and photosynthetic capacity under drought stress in citrus.

### Photosynthetic pigment dynamics in citrus leaves under drought and microbial symbiosis

Water deficit markedly diminished Chlorophyll content in citrus, with non-symbiotic stressed plants (D) displayed reductions in Chl *a*, Chl *b*, and total chlorophyll of approximately 14%, 35%, and 21%, respectively, compared to controls treatment (Fig. 4A-C).

**Fig. 4.**
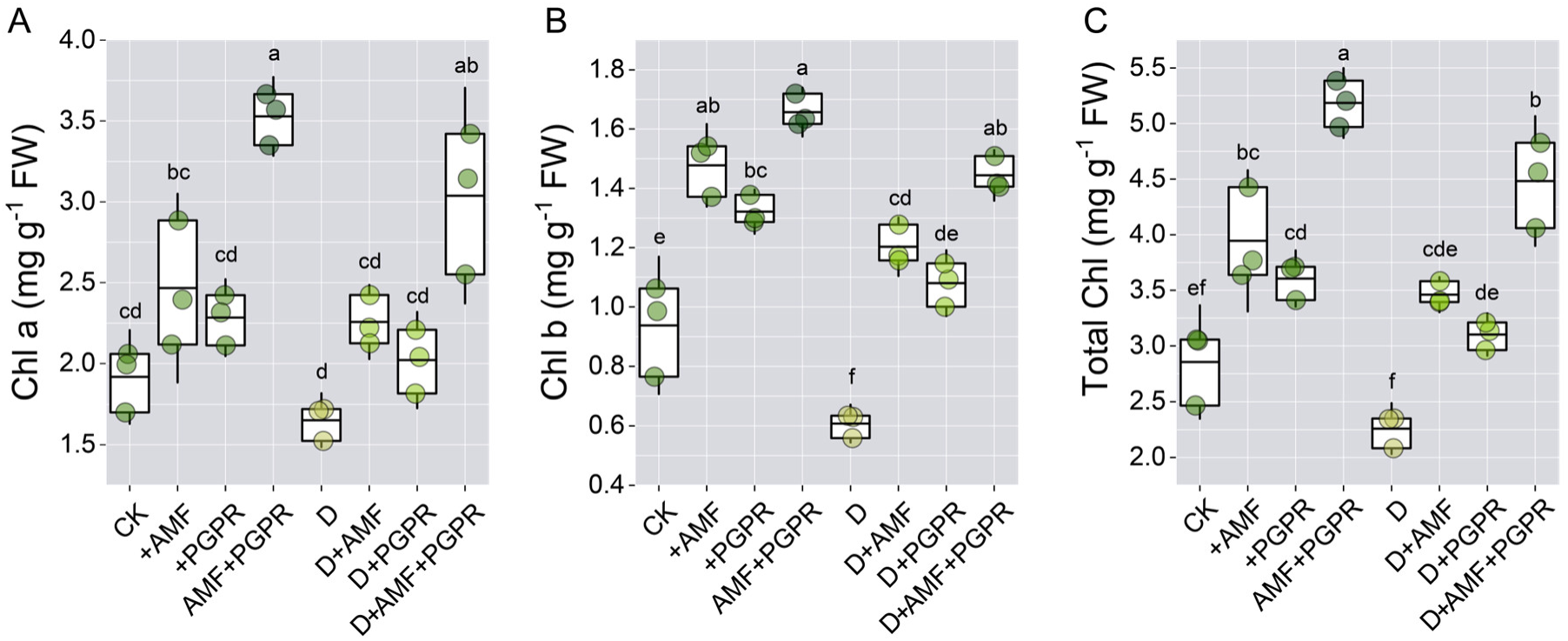
Effect of AMF and PGPR on chlorophyll content in red tangerine under drought stress. **(A)** Chlorophyll a (Chl *a*, mg g⁻¹ FW), **(B)** Chlorophyll b (Chl *b*, mg g⁻¹ FW), and **(C)** Total chlorophyll (Chl *a*+*b*, mg g⁻¹ FW) in leaves under well-watered (control, CK), AMF, PGPR, AMF+PGPR, drought (D), D+AMF, D+PGPR, and D+AMF+PGPR conditions. Data are presented as means ± SD (*n* = 3 biological replicates, with 25 plants per treatment in each replicate). Statistical significance is indicated by distinct letters (*P* < 0.05, Tukey’s HSD test following ANOVA).

Under adequate moisture, symbiotic associations generally augmented pigment contents. The AMF+PGPR co-inoculation treatment yielded the most prominent elevations, in Chl *a* by 83%, Chl *b* by 76%, and total chlorophyll by 81% relative to control treatment. Individual AMF colonization significantly enhanced chlorophyll content by Chl *a*: 28.6%, Chl *b*: 57.6%, and total chlorophyll: 38.1%, outperforming PGPR inoculation by Chl *a*: 19.1%, Chl *b*: 40.9%, and total chlorophyll: 26.2%. Both treatments showed stronger effects on Chl *b* than Chl *a*. Under drought stress, the microbial symbiosis not only mitigated chlorophyll degradation but also enhanced pigment content beyond control levels by Chl *a*: 58.4%, Chl *b*: 53.9%, and total chlorophyll: 56.9%, indicating active photosynthetic upregulation rather than mere stress protection. While both AMF and PGPR attenuated chlorophyll degradation under drought, AMF inoculation provided superior protection (28-98% recovery vs 20-78% for PGPR), particularly for Chl *b*. Overall, these outcomes illustrate the substantial capacity of microbial consortia, particularly AMF+PGPR co-inoculation, to preserve chloroplast pigment stability and mitigate drought-associated photosynthetic impairment in citrus.

### Markers of oxidative stress and antioxidant defense under drought and microbial symbiosis

Histochemical assays demonstrated elevated ROS accumulation in drought-exposed citrus leaves, as indicated by intensified DAB and NBT staining for hydrogen peroxide (H₂O₂) and superoxide (O₂⁻), respectively, in drought plants compared with controls (Fig. 5A, B). Microbial inoculation markedly attenuated staining intensity, with AMF+PGPR co-inoculated leaves displaying the lowest ROS deposition under both moisture regimes, consistent with enhanced ROS scavenging capacity (Fig. S2).

**Fig. 5.**
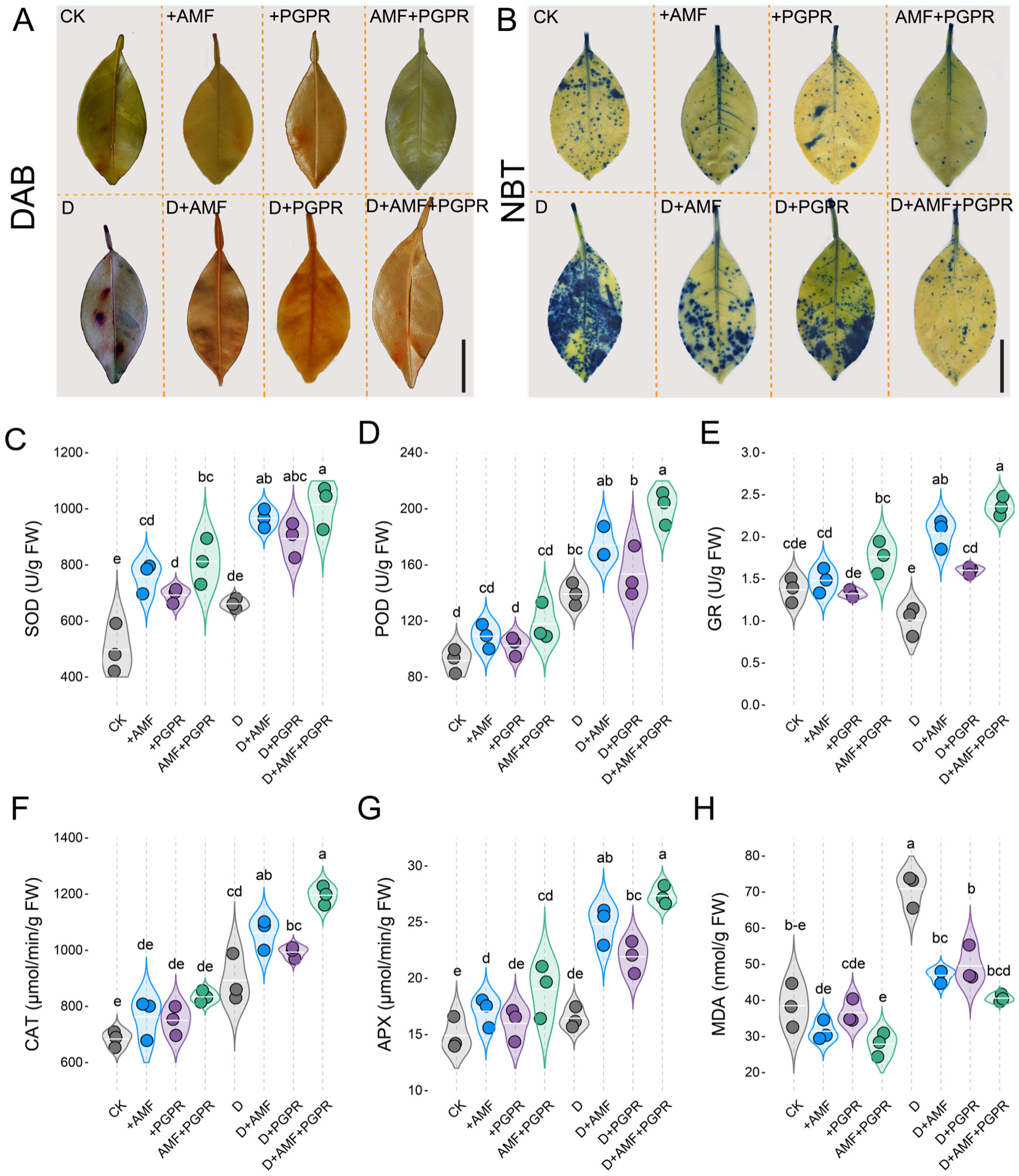
AMF and PGPR modulate oxidative stress and antioxidant responses in red tangerine under drought stress. **(A)** DAB staining of leaf tissues showing H₂O₂ accumulation under control (CK), AMF, PGPR, AMF+PGPR, drought (D), D+AMF, D+PGPR, and D+AMF+PGPR conditions. **(B)** NBT staining indicating superoxide anion (O₂⁻) accumulation across treatments. **(C-H)** Quantitative physiological parameters: **(C)** Superoxide dismutase (SOD) activity (µg min⁻¹ g⁻¹ FW), **(D)** Peroxidase (POD) activity (µg min⁻¹ g⁻¹ FW), **(E)** Glutathione reductase (µg FW), **(F)** Catalase (CAT) activity (µmol min⁻¹ g⁻¹ FW), **(G)** Ascorbate peroxidase (APX) activity (µmol min⁻¹ g⁻¹ FW), **(H)** Malondialdehyde (MDA) content (nmol g⁻¹ FW). Scale bars: 1 cm. Data are presented as mean ± SD (*n* = 3 biological replicates), and means marked with distinct letters are significantly different at *P* < 0.05 using Tukey’s HSD after ANOVA.

Enzymatic analyses further revealed that drought altered antioxidant homeostasis. Compared with control, drought increased SOD by 33.0%, while POD, GR, CAT, and APX changed by +51.6%,−26.3%, +30.5%, and +10.2%, respectively (Fig. 5C-G). Under optimal hydration, inoculated plants exhibited higher antioxidant capacity, with AMF+PGPR producing the strongest enhancements, compared with control. While SOD increased by 63.2%, POD by 28.3%, GR by 28.4%, CAT by 22.0%, and APX by 27.6%. Under drought conditions, microbial symbiosis substantially reinforced antioxidant defenses. AMF increased SOD, POD, GR, CAT, and APX by 45.9%, 25.0%, 103%, 18.9%, and 51.1% relative to drought, while PGPR increased these enzymes by 35.1%, 10.2%, 58.4%, 11.2%, and 33.4%. Notably, the AMF+PGPR treatment delivered the strongest recovery, elevating SOD by 53.4%, POD by 44.7%, GR by 133.6%, CAT by 33.9%, and APX by 66.4% compared with drought treatment, demonstrating a synergistic enhancement of cross-enzyme ROS detoxification.

Malondialdehyde (MDA), a marker of lipid peroxidation, increased by 84.0% in drought, compared with control (Fig. 5H). AMF and PGPR reduced drought-induced MDA accumulation by 33.8% and 30.1% compared with drought, while the AMF+PGPR co-inoculation achieved the greatest protection, reducing MDA by 42.7% compared with drought and maintaining it 5.4% above control. These integrated biochemical and histochemical patterns underscore the superior efficiency of AMF+PGPR co-colonization in bolstering ROS detoxification and safeguarding membrane stability during drought stress.

### Nutrient concentration in citrus tissues under drought and microbial symbiosis

Drought stress profoundly disrupted nutrient homeostasis in citrus relative to the control treatment. Specifically, leaf N declined by 32.6%, root N by 37.6%; leaf P by 23.2%, root P by 28.6%; leaf K by 18.1%, root K by 26.5% (Fig. 6A-C). Micronutrient depletion followed analogous trends: leaf Mn decreased by 39.4%, root Mn by 31.6%; leaf Zn by 14.9%, root Zn by 26.9%; leaf Fe by 22.0%, and root Fe by 28.5% (Fig. 6D-F).

**Fig. 6.**
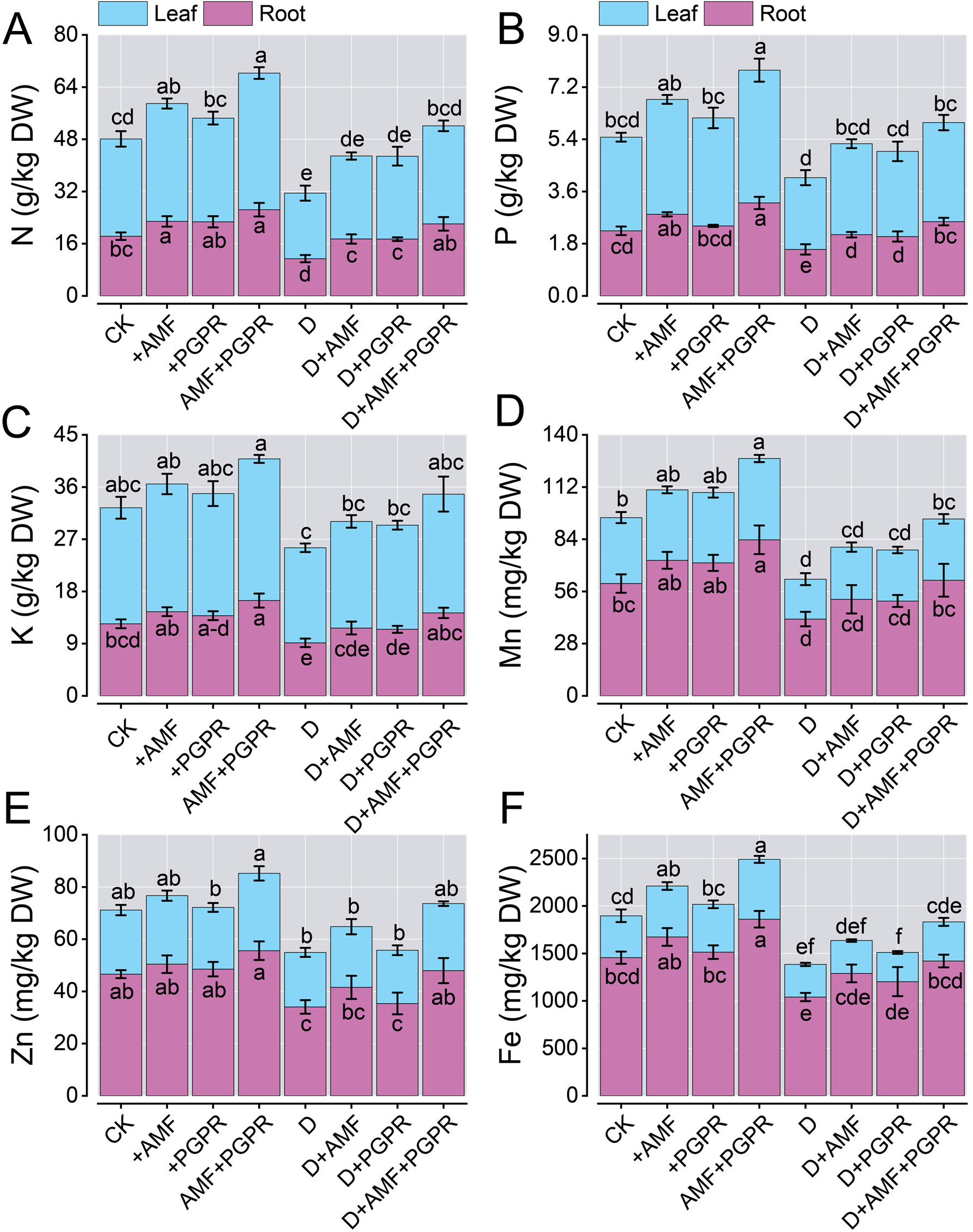
AMF and PGPR effects on nutrient accumulation in red tangerine under drought stress. Concentrations of **(A)** Nitrogen (N), **(B)** Phosphorus (P), **(C)** Potassium (K), **(D)** Manganese (Mn), **(E)** Zinc (Zn), and **(F)** Iron (Fe) in leaf (blue) and root (purple) tissues (g or mg kg⁻¹ DW) under well-watered (control, CK), AMF, PGPR, AMF+PGPR, drought (D), D+AMF, D+PGPR, and D+AMF+PGPR conditions. Data are presented as mean ± SD (*n* = 3 biological replicates), and means marked with distinct letters are significantly different at *P* < 0.05 using Tukey’s HSD after ANOVA.

Microbial inoculation, particularly the AMF+PGPR consortium, robustly reversed these deficits. Under well-watered conditions, dual inoculation elevated leaf N by 40.4%, P by 41.8%, K by 22.1%, Mn by 23.2%, Zn by 20.7%, and Fe by 43.1% over CK, with root increases of 44.4% (N), 42.9% (P), 32.3% (K), 39.0% (Mn), 19.3% (Zn), and 27.8% (Fe). Individual AMF and PGPR treatments showed moderate improvements (4-25% across elements), with AMF generally surpassing PGPR in root nutrient acquisition. Under drought, AMF+PGPR not only restored but often exceeded control levels, increasing leaf N, P, K, Mn, Zn, and Fe by 0.8%, 5.6%, 2.2%, 7.2%, 4.5%, and 6.7%, respectively, relative to CK, while root values increased by 20.6% (N, \**p*\*<0.01), 14.2% (P, \**p*\*<0.05), 15.2% (K, \**p*\*<0.05), 3.0% (Mn, ns), 2.9% (Zn, ns), and 2.4% (Fe, ns). Solitary AMF and PGPR provided partial mitigation (5-30% recovery vs D), highlighting the consortium’s superior capacity to enhance nutrient solubility, translocation, and osmotic resilience. These patterns affirm the pivotal function of AMF+PGPR co-colonization in facilitating nutrient acquisition and homeostasis amid drought in citrus.

### Phytohormone concentrations in citrus leaves under drought and microbial symbiosis

Drought stress significantly altered the endogenous phytohormone profile in citrus leaves. Drought-stressed plants (D) exhibited higher levels of ABA (+39.4%), ACC (+61.7%), IAA (+28.9%), and JA (+130.7%) compared with control, while reductions were observed in *t*ZR (−31.7%) and GA1 (−28.5%) (Fig. 7A-F).

**Fig. 7.**
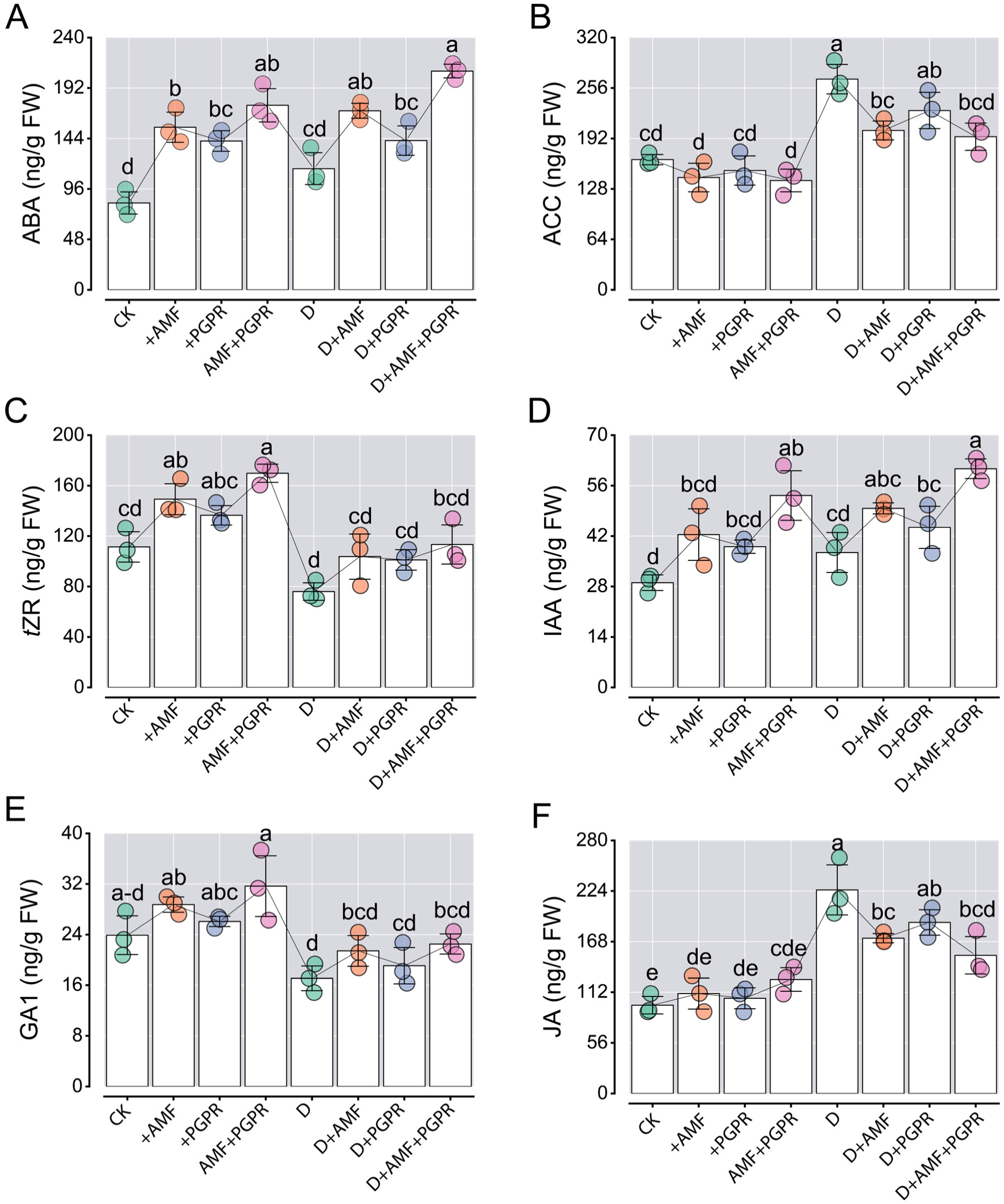
Effects of AMF and PGPR phytohormone levels in red tangerine leaves under drought stress. Concentrations of phytohormones in leaf tissues: **(A)** Abscisic acid (ABA), **(B)** 1-Aminocyclopropane-1-carboxylic acid (ACC), **(C)** trans-Zeatin riboside (tZR), **(D)** Indole-3-acetic acid (IAA), **(E)** Gibberellic acid (GA₁), and **(F)** Jasmonic acid (JA) under well-watered (control, CK), AMF, PGPR, AMF+PGPR, drought (D), D+AMF, D+PGPR, and D+AMF+PGPR conditions. Data are presented as means ± SD (*n* = 3 biological replicates, with 15 plants per treatment in each replicate). means marked with distinct letters are significantly different at *P* < 0.05 using Tukey’s HSD after ANOVA.

Microbial inoculation augmented phytohormone biosynthesis under well-watered conditions. The combined AMF+PGPR treatment induced the most significant increases, with ABA (+112.2%), *t*ZR (+52.5%), IAA (+83.3%), GA1 (+32.5%), and JA (+29.2%) higher than CK, while ACC was reduced by 16.0%. Individually, AMF inoculation led to moderate increases (ABA: +87.0%, *t*ZR: +34.0%, IAA: +46.0%, GA1: +20.2%, JA: +13.2%), with a slight decline in ACC (−13.8%), outperforming PGPR (ABA: +71.1%, *t*ZR: +22.5%, IAA: +34.4%, GA1: +9.1%, JA: +7.9%; ACC: −8.4%). Under drought stress, microbial symbiosis mitigated the hormonal imbalances induced by drought conditions. AMF+PGPR co-inoculation was the most effective, increasing ABA (+80.5%), *t*ZR (+49.1%), IAA (+62.1%), and GA1 (+31.7%) compared to D, while reducing ACC (−27.3%) and JA (−32.2%). Individual treatments with AMF and PGPR provided lesser, yet significant improvements: AMF increased ABA (+47.7%), *t*ZR (+36.4%), IAA (+32.8%), and GA1 (+25.3%), with reductions in ACC (−24.3%) and JA (−23.7%); PGPR increased ABA (+23.1%), *t*ZR (+33.0%), IAA (+18.6%), and GA1 (+11.7%), with decreases in ACC (−14.9%) and JA (−16.0%). These results demonstrate the combined effect of AMF and PGPR co-inoculation on hormone homeostasis and promoting drought resilience in citrus.

### Overall transcriptomic reprogramming in citrus leaves under drought and microbial treatments

Following adapter removal and quality filtering, 1,070.6 M clean reads were generated across the 24 cDNA libraries, averaging 44.6 M per library with Q30 ≥ 95.9% (Tab. S1). No significant variation in read numbers was detected between inoculated and uninoculated plants or between well-watered and drought treatments. The average total and unique mapping ratios were 94.7% and 96.7%, respectively. Principal component analysis (PCA) and hierarchical clustering heatmap demonstrated strong reproducibility among biological replicates, with no apparent outlier. The most significant changes in gene expression in citrus leaves were observed in the CK vs. D group (Fig. 8A, B). Our research revealed a large number of DEGs in the CK vs. D, CK vs. +AMF, CK vs. +PGPR, D vs. D+AMF, D vs. D+PGPR, CK vs. AMF+PGPR, D vs. D+AMF+PGPR, +AMF vs. +PGPR, and D+AMF vs. D+PGPR groups. PCA analysis revealed that different treatments under drought stress and microbial treatments were grouped in distinct regions of the plot, indicating that the gene expression profiles of the samples were consistent. PCA of RNA-seq data disclosed robust segregation of transcriptomes, with PC1 capturing 73% of variance and primarily delineating drought-stressed from well-watered samples, while PC2 (14.7%) highlighted microbial inoculation effects (Fig. 8A).

**Fig. 8.**
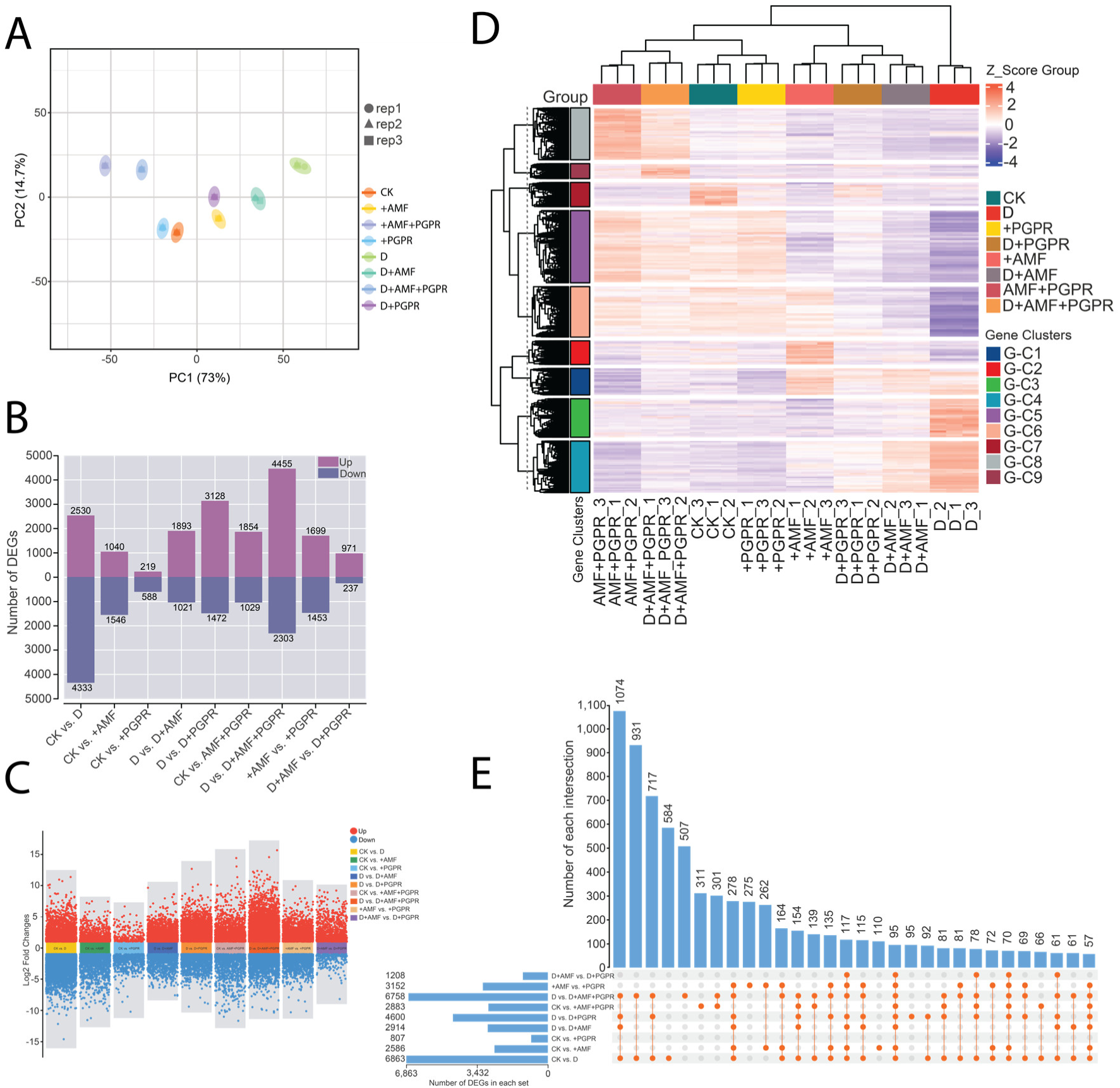
The distribution of Differentially expressed genes (DEGs) under microbial symbiosis and drought stress of citrus leaves after 70 days. **(A)** Principal Component Analysis (PCA) score plot depicting the sample distribution with PC1 and PC2contributing 73% and 14% of the variance, respectively, after treatments for 70 days. (**B)** Bar plot depicting the number of DEGs in nine comparison groups with upregulated (purple) and downregulated (blue) genes indicated among different treatments in citrus leaves. (**C)** Complex volcano plot displaying the upregulated (red) and downregulated (blue) genes in nine pairwise comparison groups. (**D)** Cluster heatmap of dynamic gene expression showing expression patterns (Z-score scaled) grouped by treatments and gene clusters. **(E)** UpSet plots showing the overlapping and uniquely associated DEGs among various groups.

Differential gene expression analysis identified substantial transcriptional shifts across pairwise comparisons, with totals 807 (CK vs. +PGPR), 6863 (CK vs. D) and 6758 (D vs. D+AMF+PGPR), the latter reflecting pronounced amelioration by combined inoculation under stress (Fig. 8B and Tab. S3). Up-regulated genes predominated in most contrasts, such as 4455 in D vs. D+AMF+PGPR and 3128 in D vs. D+PGPR, indicating activation of tolerance mechanisms. Multi-volcano plots depicted log2 fold changes spanning ±15, with dense clusters of significantly altered genes (P < 0.05, |log2 FC| > 1) in drought-related comparisons (Fig. 8C). Hierarchical clustering partitioned DEGs into nine groups exhibiting distinct profiles, including clusters upregulated in inoculated stressed samples (e.g., G-C3, G-C7) and downregulated under drought alone (e.g., G-C2, G-C5), suggestive of coordinated regulation in stress mitigation (Fig. 8D, Tab. S4). Upset analysis revealed extensive overlaps among DEG sets, with AMF+PGPR application under drought decreased the uniquely present DEGs 1074 to 931 alongside intersections in key pairs like CK vs. D and D vs. D+AMF+PGPR containing 584 and 507 gene, underscoring shared and treatment-specific pathways (Fig. 8E). Collectively, these patterns illustrate profound drought-induced perturbations and the synergistic transcriptional modulation by AMF+PGPR, likely underpinning enhanced resilience.

### GO and KEGG enrichment analyses of the differentially expressed genes

We observed significant Gene Ontology (GO) enrichment among DEGs in pairwise treatment comparisons, indicating prominent changes in biological processes (BP), cellular components (CC), and molecular functions (MF) linked to abiotic stress responses and cellular homeostasis (Fig. S3A, Tab. S5). Predominant BP categories included response to stimulus (GO:0050896), response to stress (GO:0006950), and cell wall organization or biogenesis (GO:0071554), response to external stimulus (GO:0009605), and cutin biosynthetic process (GO:0010143). These were most evident under drought conditions (CK vs. D), signifying amplified activation of protective pathways for structural remodeling and environmental adaptation. CC terms predominantly featured membrane (GO:0016020), plasma membrane (GO:0005886), cell periphery (GO:0071944), and external encapsulating structure (GO:0030312), suggesting reinforced cell wall integrity and organelle involvement during stress period. MF annotations highlighted catalytic activity (GO:0003824), transcription regulator activity (GO:0140110), DNA-binding transcription factor activity (GO:0003700), oxidoreductase activity (GO:0016491), antioxidant activity (GO:0016209), and cellulase activity (GO:0008810), indicating enhanced enzymatic adjustments for carbohydrate metabolism, redox balance, and motility. The reduced enrichment of stress-associated terms in AMF or PGPR treated plants under drought suggests improved stress acclimation through moderated metabolic and defense responses. Nonetheless, consistent overrepresentation of cytoskeletal motor activities and kinesin-related components across multiple comparisons suggests a potential contribution of microtubule dynamics to plant stress tolerance.

The Kyoto Encyclopedia of Genes and Genomes (KEGG) pathway analysis further elucidated metabolic reprogramming, with the top 24 pathways centering on carbohydrate, amino acid, and secondary metabolite biosynthesis (Fig. S3B). Glyoxylate and dicarboxylate metabolism emerged as highly significant (low adjusted P-value, Padj) in drought-exposed samples (CK vs. D), facilitating carbon recycling and energy conservation amid water deficit. In response to drought stress, several key pathways were enriched, with photosynthesis playing a central role in both drought and microbial treatments, highlighting its importance in drought and microbial responses. Enrichment of photosynthesis-antenna proteins in drought treatments with AMF and PGPR suggested enhanced light capture under microbial symbiosis. Starch and sucrose metabolism was notably enriched in D vs. D+PGPR and D vs. D+AMF+PGPR, indicating altered carbohydrate partitioning. Phenylpropanoid biosynthesis, crucial for lignin and flavonoid production in stress tolerance, was universally enriched across all comparisons (Fig. S4). Additionally, plant hormone signaling, particularly involving auxin, abscisic acid, and cytokinins, was enriched in drought conditions, suggesting that hormonal regulation plays a role in stress and growth. Galactose metabolism, which is involved in cell wall modification, was enriched under drought conditions with AMF and PGPR treatments. Cutin, suberin, and wax biosynthesis, which aids in water conservation, was enriched in drought treatments with AMF+PGPR, reflecting enhanced cuticle formation. These results indicate common metabolic reprogramming under drought and microbial treatments, with photosynthesis, phenylpropanoids, and water retention pathways being central to stress adaptation. These findings demonstrate that drought and microbial treatments coordinately modulate photosynthesis, carbohydrate metabolism, secondary metabolite synthesis, hormonal signaling, and cuticle biogenesis in citrus rootstocks. The combined effects of AMF and PGPR enhance drought tolerance through integrated metabolic and physiological adjustments.

### Identification of potential hub genes related to drought stress and microbial responses using a weighted gene co-expression network (WGCNA)

To explore the contribution of transcription factors (TFs) to drought responses in citrus rootstock and their regulation by AMF and PGPR, WGCNA was employed to detect important gene modules and hub genes related to drought responsive traits. The analysis utilized the FPKM expression dataset together with phenotypic and physiological parameters.

WGCNA analysis clustered all expressed genes into different color-coded co-expression modules via hierarchical clustering (Fig. 9A), with the turquoise module showing strong correlations with core drought resilience traits of citrus rootstocks, indicating their potential roles in mediating the boosted physiological performance of AMF-PGPR co-inoculated plants under drought stress; furthermore, the topological overlap matrix (TOM) heatmap (Fig. 9B) revealed gene co-expression effectiveness across modules, and the turquoise module showed notably high intramodular connectivity, reflecting synchronized transcriptional regulatory cascades that underpin adaptive drought responses in beneficial microbe-inoculated citrus rootstocks. Module-trait correlation analysis revealed that several modules’ eigengenes were associated with specific physiological traits and treatment conditions subjected to AMF-PGPR symbiosis and drought stress. Consistent with its functional relevance, the MEturquoise module contained the largest number of genes (1698), followed by blue (1230 genes) and brown (734 genes) (Fig. 9D, Tab. S6). Among the MEturquoise module, 1275 genes were positively correlated with physiological traits such as stomatal conductance Gs (*r*^2^=0.62, *P*=6E-181), N-R (*r*^2^=0.68, *P*=1E-200), shoot FW (*r*^2^=0.62, *P*=6.6E-181), Root FW (*r*^2^=0.64, *P*=2.6E-196), and N-L (*r*^2^=0.61, *P*=1.3E-173), respectively, showing significant positive associations. Particularly, the MEturquoise module demonstrated a strong positive correlation with microbial inoculation and drought stress (Fig. 9C), while these traits exhibited negative correlations with some modules. In contrast, MDA (*r*^2^=0.67, *P*=1E-200) showed a negative correlation with MDA (Tab. S7).

**Fig. 9.**
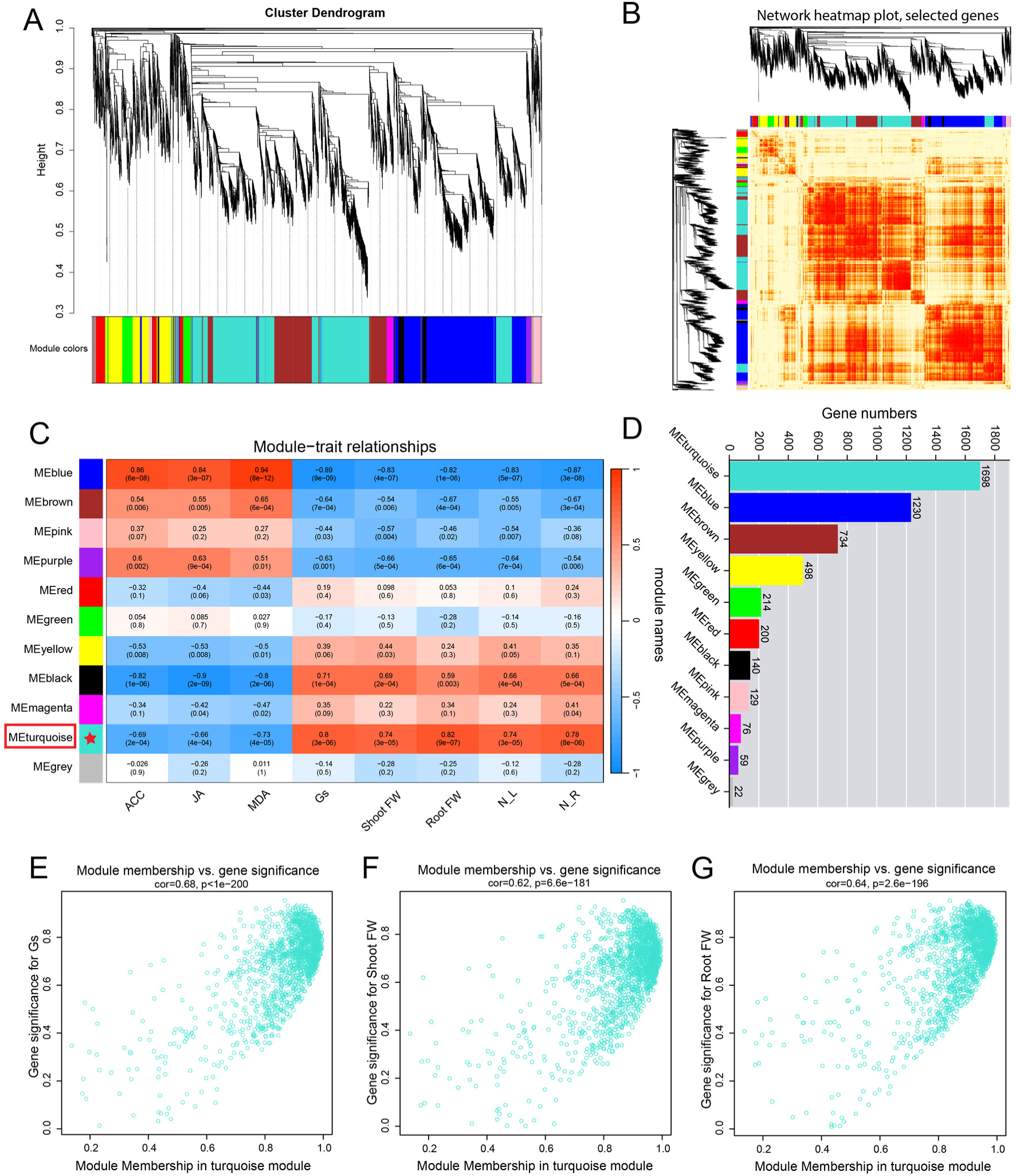
Weighted gene co-expression network analysis (WGCNA) of DEGs in citrus leaves under microbial and drought stress. (**A**) Hierarchical clustering dendrogram of co-expressed modules. (**B**) Heatmap depicting topological overlap matrix among selected genes, illustrating network connectivity. (**C**) Correlation heatmap between module eigengenes and physiological traits. (**D**) Bar plot showing the number of genes assigned to each module. (**E-G**) Scatter plots of module membership versus gene significance in the turquoise module for (**E**) stomatal conductance, (**F**) shoot fresh weight, and (**G**) root fresh weight.

Moreover, we detected the relationships between module membership and gene significance under drought stress in the MEturquoise module. Genes within the MEturquoise had a positive correlation among module membership and gene significance for Gs, shoot FW, and root FW (Fig. 9E-G), highlighting its role in stress-responsive mechanisms. These results imply that the MEturquoise module contains genes substantial for regulating plant growth and development during drought in citrus, making it a key candidate for further investigation.

To identify key regulators of the citrus response to drought and microbial inoculation, we focused on the MEturquoise module, which exhibited the strongest correlation with our physiological traits. This module with connectivity (*K _within_*) values ranging from 3.12 to 907.85. Using a stringency threshold of the top 20% *K _within_* values, we identified the primary hub genes of the network. Within this high-connectivity group, iTAK analysis classified 18 genes as Transcription Factors (TFs), representing families such as MYB, MYB-related, and NF-YB (Fig. 10A, Tab.S8).

**Fig 10.**
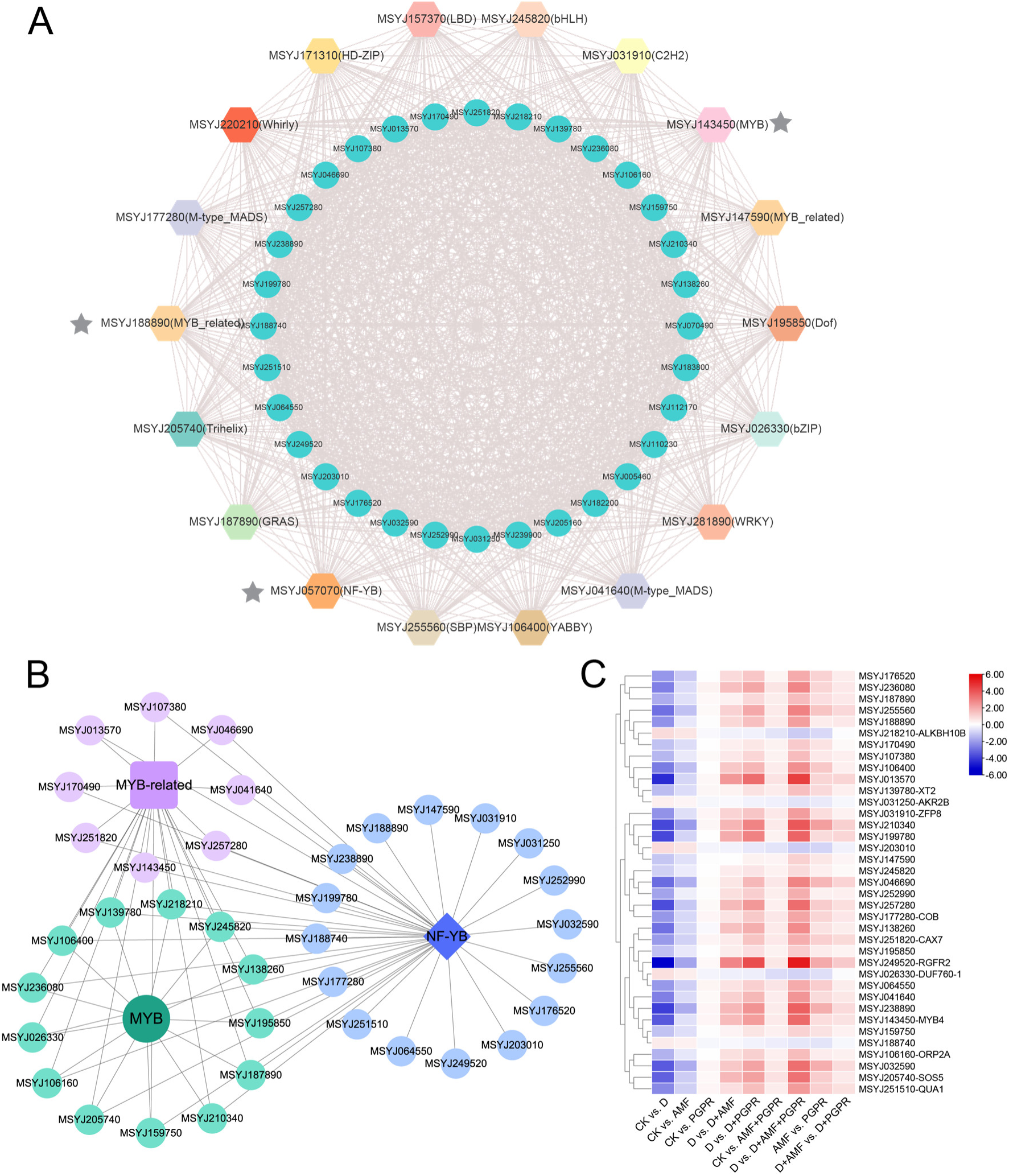
Transcription factor, gene interaction networks and expression profiles of drought-responsive hub genes in citrus. (**A**) Co-expression network of differentially expressed genes (DEGs) and transcription factors (TFs), highlighting key TF families including MYB, bHLH, WRKY, bZIP, NF-YB, and MADS. The network illustrates the extensive regulatory connectivity between TFs (colored nodes) and their associated target genes (blue nodes). (**B**) Subnetwork illustration highlighting the MYB, MYB-related, and NF-YB transcription factors and their interacting hub genes. (**C**) Heatmap of representative hub genes showing transcriptional changes across treatments, with color gradients indicating relative expression (red = upregulated; blue = downregulated).

From this regulatory framework, we selected three primary TFs as central hubs based on their peak *K _within_* values: *CrMYB4* (MSYJ143450; *K _within_* = 866.19), *CrMYB-related* (MSYJ188890; *K _within_* = 868.70), and *CrNF-YB* (MSYJ057070; *K _within_* = 877.78). These genes are orthologous to *Arabidopsis AtMYB4*, *AtMYB-related*, and *AtNF-YB*, respectively. Sub-network analysis revealed distinct co-expression patterns: the *CrMYB4* network comprised 13 co-expressed genes, the *CrMYB-related* network contained 8 genes, and the *CrNF-YB* network included 16 genes (Fig. 10B).

To elucidate the functional relevance of these networks, we isolated 12 critical hub genes that exhibited significant differential expression across treatments (Fig. 10C). These included five MYB members (e.g., *CrSOS5* and *CrORP2A*), two MYB-related genes (*CrMYB4* and *CrCAX7*), and five NF-YB members (*CrCOB*, *CrQUA1*, and *CrZFP8*). Orthology-based functional annotation via TAIR BLAST suggested these hubs are integral to growth modulation and signaling. For example, *CrZFP8* and *CrMYB4* are implicated in stress-responsive growth regulation, while *CrQUA1* and *CrRGFR2* are associated with calcium-mediated signaling and root architecture development. Notably, the expression of *CrRGFR2*, *CrSOS5*, *CrMYB4*, *CrQUA1*, and *CrZFP8* was markedly upregulated in the D+AMF+PGPR group compared to drought-stressed plants (D), whereas these same genes were suppressed under drought conditions alone (CK vs. D). This expression reversal suggests a “molecular switch” mechanism, where microbial inoculation restores or enhances the expression of key developmental and stress-signaling genes to mitigate drought-induced damage (Fig. 10C).

### qRT-PCR verification of genes

Additionally, we validated the expression profiles of 12 genes through qRT-PCR, confirming that their relative expression levels (Fig. S6) largely aligned with the RNA-seq results. A significant correlation was observed between the expression changes (log2 fold-change) from RNA-seq and qRT-PCR, further supporting the reliability of the RNA-seq data in identifying drought-responsive genes in citrus leaves treated with AMF and PGPR, either separately or in combination, under drought stress for 70 days. Notably, the expression levels of *CrRGFR2*, *CrMYB4*, and *CrZFP8* were significantly elevated in response to combined AMF and PGPR inoculation under drought conditions, while *CrDUF760-1*, *CrAKR2B*, and *CrALKBH10B* exhibited a marked reduction in expression under the same treatment (Fig. S6).

## Discussion

Drought-induced oxidative stress severely limits citrus performance because elevated ROS damages membranes and enzymes, degrades photosynthetic pigments, and impairs light capture and energy conversion (Gill & Tuteja, 2010), while simultaneous inhibition of root development and nutrient uptake further depresses photosynthetic capacity and whole-plant vigor (Rao *et al*., 2024). To alleviate these constraints, growers increasingly use microbial inoculants such as PGPR and AMF to reduce mineral fertilizer inputs, improve nutrient bioavailability, and buffer plants against environmental stress (Zhyr *et al*., 2025), with beneficial taxa including nitrogen-fixing *Pseudomonas* spp. and AMF such as *F. mosseae,* enhancing mineral acquisition, water uptake, and resilience against biotic and abiotic stressors (Abhilash *et al*., 2016; Zhang *et al*., 2019). Within this context, our pot experiment with drought-stressed Citrus rootstocks showed that sole AMF, PGPR, and especially their co-inoculation reprograms plant water relations, water-use efficiency, biomass allocation, phytohormone profiles, nutrient status, oxidative stress markers, and stress-responsive gene expression, thereby mitigating drought-induced growth suppression. It is consistent with previous work on AMF–bacteria consortia and indigenous microbiomes (Trivedi *et al*., 2020), and the established role of *Pseudomonas* strains as mycorrhiza helper bacteria that promote AMF colonization (Frey-Klett *et al*., 2007; Cosme & Wurst, 2013; Ain *et al*., 2024) together with the reciprocal stimulation of *Pseudomonas* growth by AM fungi (Zhang *et al*., 2024).

### Plant growth and health

Improvement in plant growth and water relations serves as an early indicator of stress resilience, often attributed to phytohormone modulation and enhanced nutrient acquisition via microbial inoculants (Chen *et al*., 2025). In this study, AMF and PGPR significantly improved multiple shoot and root traits under both well-watered and drought conditions, with the AMF+PGPR combination increased height (43.4%) and biomass (51.8%) relative to non-inoculated drought plants (Fig. 1). Under drought stress, non-inoculated plants experienced significant decline in relative water content (RWC), leaf area, height, and biomass (Morade *et al*., 2025; Zeng *et al*., 2025), with RWC loss indicating reduced water retention capacity (Abdelaal *et al*., 2024). Inoculation with AMF and PGPR, particularly their co-inoculation, substantially restored RWC, preserved shoot architecture, and maintained biomass, aligning with previous findings in other species (Zou *et al*., 2017; Papadopoulou *et al*., 2022).

Root system development is critical for citrus to acquire water and nutrients under drought, with drought-induced limitations on root growth and nutrient uptake impairing photosynthesis, respiration, and cell division, thus reducing biomass production (Rao *et al*., 2024). *Pseudomonas* strains are recognised as mycorrhiza helper bacteria, promoting colonisation of both ectomycorrhizae and AMF in various studies (Li *et al*., 2023). In our study, drought treatment reduces mycorrhizal colonisation, though inoculated plants show improved biomass production, root: shoot ratios, and root morphology under drought conditions compared to uninoculated plants (Fig. 2). In contrast, non-AMF-treated plants lack mycorrhizal colonisation. AMF-inoculated plants, especially with PGPR, exhibit significant root colonisation. This study confirms that drought stress limits *F. mosseae* colonisation, in line with the previous studies on citrus (Zhang *et al*., 2020; Zheng *et al*., 2024). In our study, drought impaired root biomass and architecture, but microbial inoculation, particularly AMF+PGPR co-inoculation, significantly enhanced root biomass and length, with up to a 57% improvement under drought conditions (Fig. 2). This indicates that the combined effects of AMF-driven root proliferation and PGPR-mediated nutrient solubilization and hormone production are key mechanisms for citrus drought adaptation. These findings align with similar results in tobacco, maize, and wheat, suggesting the broad potential of AMF–PGPR consortia to enhance drought resilience in cropping systems under increasing aridity (Rehman *et al*., 2025; Zeng *et al*., 2025).

### Improved photosynthetic performance

Stomatal regulation is central to drought responses because reductions in stomatal density and aperture rapidly restrict CO₂ influx and drive the decline in photosynthetic rate under water deficit (Augé *et al*., 2015). Consistent with this, SEM showed that drought-stressed plants had fewer stomata with narrower pores, whereas microbial inoculation, especially the AMF+PGPR consortium, largely preserved stomatal density and aperture and increased stomatal size by ∼27% versus controls (Fig. 3). AMF and PGPR enhance photosynthesis through effects on stomatal conductance, hormone balance and root-mediated water and nutrient uptake (Khan *et al*., 2024; Abrar *et al*., 2025). Accordingly, drought sharply reduced Pn, Tr, Ci and Gs in our study, but co-inoculation substantially reversed these losses and even raised Gs above control values (Fig. 3). These findings indicate that AMF+PGPR maintain plant water status and stomatal behaviour via coordinated improvements in root function and hormonal signalling, thereby preserving internal CO₂ and sustaining carbon assimilation during drought (Li *et al*., 2025; Zou *et al*., 2025). Collectively, the data show that AMF+PGPR consortia stabilise gas exchange and photosynthetic performance under water deficit, providing a mechanistic basis for improved drought-time productivity and agreeing with prior reports (Khan *et al*., 2024; Zheng *et al*., 2024; Zou *et al*., 2025).

In our experiment, drought markedly decreased chlorophyll, particularly Chl b, and constrained biomass accumulation; by contrast, AMF+PGPR co-inoculation not only offset these losses but raised Chl *a* and Chl *b* by 58.4% and 53.9%, respectively (Fig. 4). Similar pigment increases after AMF or AMF-bacterial consortia in trifoliate orange, *Arabidopsis* and other crops support this interpretation (He *et al*., 2020; Yang *et al*., 2021; Nader *et al*., 2024), and underscore that well-chosen bioinoculants can preserve light harvesting and carbon gain, and thus sustain growth, in drought-prone systems (Khan *et al*., 2024; Zeng *et al*., 2025).

### Oxidative stress and antioxidant defense

Drought altered the balance of core ROS-scavenging systems in a trait-specific manner rather than producing a simple generalized suppression in citrus. Drought stress leads to an increase in ROS such as H_2_O_2_, and O_2_^-^, which cause oxidative damage in plant cells, including lipid peroxidation, protein degradation, and pigment bleaching (He *et al*., 2021). AMF and PGPR improve the plant’s resilience to damage by upregulating its antioxidant defense mechanism. Inoculation with AMF improves drought resilience by elevating antioxidant enzyme activity, lowering ROS levels, and enhancing water uptake through fungal hyphae, thus reducing the generation of ROS (Huang *et al*., 2017). Similarly, PGPR inoculation enhances the actions of antioxidant enzymes, for example, CAT and SOD, safeguarding plant cells against drought-induced oxidative stress. (Uzma *et al*., 2022). The combination of AMF and PGPR can synergistically reduce ROS levels and improve plant performance by enhancing oxidative stress tolerance, preserving cell membrane integrity, and boosting photosynthetic efficiency under water deficit conditions (Ben Laouane *et al*., 2019) (Ali *et al*., 2022). In this study, SOD activity increased ∼33.0% under drought compared with controls, yet strong NBT staining persisted in drought, indicating that enhanced superoxide production outpaced primary dismutation and that spatial ROS accumulation remained pronounced (Noctor & Foyer, 1998). By contrast, GR was markedly decreased-26.3% by drought, signifying compromised glutathione recycling and a bottleneck in the ascorbate–glutathione cycle, POD, CAT and APX increase ∼51.6%, ∼30.5% and∼10.2% by drought, respectively (Fig. 5), suggesting a partial, and ultimately insufficient, upregulation of enzymatic H₂O₂ turnover pathways. This mixed response differs from many reports in which drought alone elicits coordinated upregulation of the full suite of antioxidants or where AMF and/or PGPR enhance such responses (Brilli *et al*., 2019; Liu, Y *et al*., 2023). Crucially, AMF+PGPR co-inoculation more effectively restructured the antioxidant network under water deficit than alone. relative to droughted controls, the consortium elevated GR, SOD, POD, CAT, and APX, thereby restoring glutathione recycling, accelerating H₂O₂ turnover, and stabilising chloroplastic redox poise. Single inoculations offered intermediate recovery, consistent with additive but non-identical modes of action between fungal and bacterial partners (He *et al*., 2020; Mekureyaw *et al*., 2022). MDA integrated these biochemical outcomes at the level of membrane integrity: drought increased MDA versus control, whereas AMF and PGPR individually reduced drought-induced MDA respectively, and AMF+PGPR lowered MDA relative to drought, leaving values only ∼5.4% above well-watered plants (Fig. 5). These findings align with the previous reports on citrus, and tobacco showing reduced lipid peroxidation applying microbial inoculation (Huang *et al*., 2017; Sadeghi *et al*., 2020; Begum *et al*., 2022). These indicate that AMF–PGPR consortia do not merely add protective effects but reorganise antioxidant fluxes to preserve membrane stability and enhance drought resilience in citrus.

### Drought effect on nutrients

Plants exposed to drought showed marked declines in leaf and root N, P, K, Mn, Zn and Fe, indicating concurrent limitations to photosynthesis, protein synthesis and osmotic and redox regulation (Sheshbahreh *et al*., 2019; De Vries *et al*., 2020; Khare *et al*., 2025; Sulaman *et al*., 2025). AMF and PGPR offer a complementary, sustainable approach to mitigate nutrient losses by enlarging the absorptive interface, mobilizing inaccessible nutrient pools, and modifying rhizosphere biochemistry and biodiversity to favor nutrient release and uptake (Sheshbahreh *et al*., 2019; De Vries *et al*., 2020). Rhizospheric bacteria such as *Pseudomonas* act as mycorrhizal helpers by improving colonization and hyphal branching and by solubilizing N, P, K, and trace elements, thereby improving nutrient availability for the host plant (Frey-Klett *et al*., 2007; Berrios *et al*., 2023; Ain *et al*., 2024; Duan *et al*., 2024). AMF hyphae extend into soil micropores and interact with soil microbiota to accelerate organic matter turnover and nutrient exchange (Zhang *et al*., 2016; Zhang *et al*., 2018; Hestrin *et al*., 2019). Co-inoculation with AMF and PGPR consistently improves N, P, and K acquisition, reduces oxidative stress, and enhances drought resistance (Sagar *et al*., 2021; Abrar *et al*., 2025). In this study, drought reduced N, P, K, Mn, Zn and Fe in leaf and root, but the AMF+PGPR consortium reshaped this profile (Fig. 6), consistent with AMF access to distal nutrient pools and improved N and P status across hosts (Ahmed *et al*., 2025; Mazumder *et al*., 2025). The improvement of K implies microbial reinforcement of osmotic adjustment, stomatal function, and membrane stability, aligning with the role of K in antioxidant defence and reports that mycorrhizae and biofertilizers boost K uptake and drought resilience (García-Martí *et al*., 2019; Abrar *et al*., 2025). Mn, Zn, and Fe improvement under co-inoculation likely reflects AMF-mediated changes in solubility and compartmentation together with microbial phosphatase and organic-acid release, siderophore production, and transporter activation that enhance trace-element availability and sustain photosynthetic and antioxidant metabolism under stress (Rahman *et al*., 2020; Bhantana *et al*., 2021; Pan *et al*., 2023; Arifuzzaman *et al*., 2024). Overall, AMF+PGPR co-colonization reconfigures nutrient fluxes by stabilizing photosynthetic performance, antioxidant capacity, and providing a mechanistic basis for microbe-mediated drought tolerance in citrus.

### Phytohormone regulation and stress tolerance

The regulation of key physiological responses, including stomatal closure, root architecture, and growth maintenance under drought, is mediated by complex phytohormonal networks (Mathur & Roy, 2021). Inoculation with both AMF and PGPR alters phytohormone levels by increasing indole-3-acetic acid (IAA) and gibberellic acid (GA), while reducing ABA and 1-aminocyclopropane-1-carboxylate (ACC), mitigating the negative effects of drought, such as reduced stomatal conductance, senescence, and root inhibition (Chandran *et al*., 2021; Rehman *et al*., 2024). PGPRs reduce ethylene accumulation through the production of ACC deaminase, thereby alleviating stress-induced growth inhibition (Glick, 2014). Furthermore, AMF hormone distribution within the soil, strengthens plant growth and water absorption (Pérez-de-Luque *et al*., 2017; Tsukanova *et al*., 2017).

ABA is central to managing water homeostasis and enhancing stress resilience (Mathur & Roy, 2021), with AMF and PGPR, influence its levels. As shown in Figure 7, AMF and PGPR, both individually and in combination, elevate ABA levels under drought by improving stomatal regulation and water retention, in line with previous reports (Duc *et al*., 2023; Savastano & Bais, 2024). Cytokinins, especially trans-zeatin riboside (tZR), are key in regulating cell growth and water-use efficiency, with microbial symbiosis helping restore tZR levels after drought-induced reductions. AMF+PGPR co-inoculation increased tZR to support shoot growth and delaying senescence (Liu *et al*., 2013). In contrast, ACC accumulation was mitigated by both AMF and PGPR treatments, with the AMF+PGPR consortium showing the greatest suppression of ACC, preserving root activity and photosynthesis (Glick, 2004; Sati *et al*., 2023). JA increased significantly under drought, reflecting a growth-limiting defense response; however, microbial symbiosis attenuated this surge, particularly the AMF+PGPR combination, which reduced JA, helping conserve metabolic resources and maintain growth (Forchetti *et al*., 2007; Castillo *et al*., 2013). Similarly, IAA was elevated by AMF+PGPR co-inoculation, promoting root expansion and improving water uptake (Liu *et al*., 2016; Chen *et al*., 2020). Lastly, GA biosynthesis, reduced under drought, was restored by microbial treatments, with co-inoculation leading to the most substantial recovery, suggesting a mechanism for promoting shoot expansion and carbon assimilation during water stress (Sandhya *et al*., 2017). Overall, our findings underscore the synergistic potential of microbial consortia in modulating hormonal pathways to enhance drought resilience in plants.

### Microbe-modulated transcriptome reprogramming under drought

Drought rapidly engages conserved signaling networks that elevate ROS as systemic stress signals and reprogram whole-plant physiology. Plant-associated microbes intervene in these responses by mediating osmoregulation and by supplying osmolytes and signaling compounds that bolster host osmoprotection and drought-responsive transcription (Singh *et al*., 2020). Molecular studies of multipartite legume symbioses remain sparse, yet available transcriptomes show that co-inoculation produces transcriptional states distinct from single symbioses: dual inoculation of *Glycine max* with *Bradyrhizobium diazoefficiens* and *Gigaspora rosea* elicited unique gene expression changes (Sakamoto *et al*., 2019), and similar dual-symbiosis effects on host coexpression have been reported in other legume–AMF–rhizobia systems (Palakurty *et al*., 2018; Sakamoto *et al*., 2019). Across studies, AMF typically exert a larger influence on host transcription than rhizobia, consistent with their broader roles in provisioning phosphorus, nitrogen, water and other resources (Afkhami & Stinchcombe, 2016). Our RNA-seq from citrus leaves concurs: the drought contrast (CK vs D) produced the largest set of differentially expressed genes (Fig. 8) characterized by induction of canonical stress pathways and repression of photosynthetic machinery, while inoculation (AMF, PGPR, and especially AMF+PGPR) substantially reshaped that drought signature, consistent with microbe-mediated priming that enables faster, more economical responses to water deficit (Zamioudis *et al*., 2015; Liu *et al*., 2022). Weighted network analysis highlighted drought-responsive transcription factors (MYB, MYB-related, NF-YB, WRKY, bHLH, GRAS, bZIP) as highly connected hubs and therefore promising targets for breeding or genetic manipulation (Fig. S9). Pathway enrichment further shows that photosynthesis and carbon-fixation pathways are among the most affected in CK vs D, with partial recovery under single inocula and the strongest restoration under AMF+PGPR; parallel enrichment of starch/sucrose, galactose and nucleotide-sugar metabolism indicates active rerouting of photoassimilates, matching reports that AMF and PGPR sustain photochemistry and reprogram sucrose partitioning and trehalose cycling to support osmotic adjustment and antioxidant capacity (Raheem *et al*., 2018; Xu *et al*., 2018; Singh *et al*., 2019; Liu *et al*., 2022; Zheng *et al*., 2024) via SWEET transporters and invertase/SPS nodes that determine sink–source balance (Stein & Granot, 2019; Mekureyaw *et al*., 2022; Zheng *et al*., 2024). Concurrent enrichment for cutin/wax and cell-wall remodeling terms points to epidermal barrier fortification and wall polysaccharide adjustments that stabilize water status and feedback on defense signaling (Vogt, 2010; Gao *et al*., 2015; Singh *et al*., 2019; Zheng *et al*., 2024). Finally, repeated enrichment of hormone signaling (ABA, JA/SA, ethylene, auxin, cytokinin, brassinosteroid) underscores that ABA-ROS guard-cell programs are central to drought responses, while beneficial microbes modulate hormone homeostasis and sensitivity, AMF altering ABA/GA traits and PGPR influencing host hormone profiles or releasing priming volatiles, thereby reducing stress amplitude while preserving photosynthetic and carbohydrate programs (Cho *et al*., 2011; Pinheiro & Chaves, 2011; Cho *et al*., 2012; Cho *et al*., 2013; Xu *et al*., 2022; Yasmin *et al*., 2022).

### Phenylpropanoid metabolism and MYB-centered control as a regulatory nexus

Phenylpropanoid biosynthesis emerged as a consistent, central response to drought and to drought-plus-microbe treatments, implicating lignin and flavonoid branches in ROS buffering and membrane and photosystem stabilization (Dong & Lin, 2021; Jian *et al*., 2024). WGCNA identified a turquoise module that correlated positively with stomatal conductance and fresh biomass and was enriched for MYB, MYB-related, and NF-YB transcription factors (Fig. 10), families known to govern phenylpropanoid and flavonoid programmes and to coordinate redox and guard-cell responses (Dai *et al*., 2007; Dubos *et al*., 2010). qRT-PCR validated twelve hub genes from this module, *CrMYB4*, *CrZFP8*, *CrRGFR2*, *CrSOS5*, *CrQUA1*, *CrAKR2B*, *CrALKBH10B*, *CrCAX7*, *CrCOB*, *CrDUF760-1*, *CrTSD2*, and *CrXT2*, whose expression tracked the microbe-dependent shift from damage-containment under D to growth-compatible acclimation under AMF+PGPR. Notably, *CrZFP8* was downregulated by drought but upregulated in D+AMF+PGPR, consistent with Tian et al. (2024) (Tian *et al*., 2024) who showed ZFP8 negatively regulates ABA-mediated drought responses by inhibiting ZFP8 transcriptional activity, and suggesting a role for ZFP8 in balancing stress responses when microbes are present (Tian *et al*., 2024). TSD2, implicated in cell-wall integrity and meristem function, and here represented by *CrTSD2*, likewise appears positioned to link developmental regulation and stress tolerance (Krupková *et al*., 2007). *CrMYB4* showed drought-responsive dynamics, upregulated with inoculation and repressed by drought alone, mirroring reports that the overexpression of MYB4 improves tolerance to drought, salt and cold via activation of stress pathways and osmolyte accumulation (Pasquali *et al*., 2008; Lian *et al*., 2021; Yu *et al*., 2021; Kha & Nguyen, 2024). *CrCOB* and CrSOS5 were also modulated by microbial inoculation in ways that imply adaptive cell-wall remodelling and ABA/ROS-related signalling during stress, consistent with prior descriptions of COB-mediated wall modification and SOS5 (FLA4)-dependent ABA and antioxidant functions (Dinneny *et al*., 2008; Acet & Kadıoğlu, 2020). The receptor kinase *CrRGFR2*, whose expression pattern suggests reactivation by inocula and repression by drought, may coordinate root growth and architecture through PLETHORA-linked gradients in the stem-cell niche, further linking developmental plasticity to stress adaptation (Song *et al*., 2016). Together with the sugar-and cuticle/wall-related signatures, these data support a model in which AMF+PGPR steer citrus toward a MYB-centered, phenylpropanoid-rich state that stabilizes photosynthesis, membranes and growth under water limitation (Papadopoulou *et al*., 2023; Jian *et al*., 2024; Zheng *et al*., 2024).

We interpret the leaf transcriptome as the systemic output of two superimposed programs. First, drought decreases photosynthetic capacity, promotes ABA-ROS signaling, and redirects carbon into compatible solutes and structural polymers (Pinheiro & Chaves, 2011; Abhinandan *et al*., 2018; Xu *et al*., 2022). Second, microbial partners, especially AMF+PGPR, recalibrate carbon allocation (sucrose/trehalose circuits and transporters), reinforce epidermal and wall barriers, and activate MYB/NF-YB-linked phenylpropanoid modules, thereby maintaining growth with restrained stress intensity (Dubos *et al*., 2010; Vogt, 2010; Zamioudis *et al*., 2015; Stein & Granot, 2019). The dual inoculation produced the most coherent realignment and the clearest association with gas exchange and biomass in our data, which agrees with reports that combined symbioses outperform single partners under drought (Sakamoto *et al*., 2019; Yasmin *et al*., 2022; Papadopoulou *et al*., 2023; Zheng *et al*., 2024).

Our transcriptomic analysis primarily focused on leaf tissue, offering a systemic perspective of drought stress responses, but it does not resolve root-specific processes at the symbiotic interface.

Future work should incorporate tissue-resolved transcriptomics to elucidate root-localized gene expression, particularly in relation to nutrient uptake and osmotic regulation. Additionally, perturbing key transcription factors such as *CrMYB4*, *CrRGFR2*, and *CrSOS5* will help establish causal links within the turquoise module, providing a deeper understanding of microbial modulation of drought tolerance. To better capture temporal dynamics, a joint host-microbe time-series during progressive soil drying would clarify how microbial inoculation modulates hormone signaling, sugar allocation, and phenylpropanoid production in response to drought. Finally, further studies on the coordination between AMF and PGPR in mediating drought resilience through root-to-shoot signaling will improve insights into plant adaptation and guide future bio-inoculant development.

These findings provide compelling evidence of the integral roles of plant microbiomes, highlighting the intricate interactions between plant metabolism and microbial adaptation. They offer novel mechanistic insights into the molecular basis of drought resilience in plants, paving the way for leveraging these microbes for sustainable crop improvement in variable climates (Fig. 11).

**Fig. 11.**
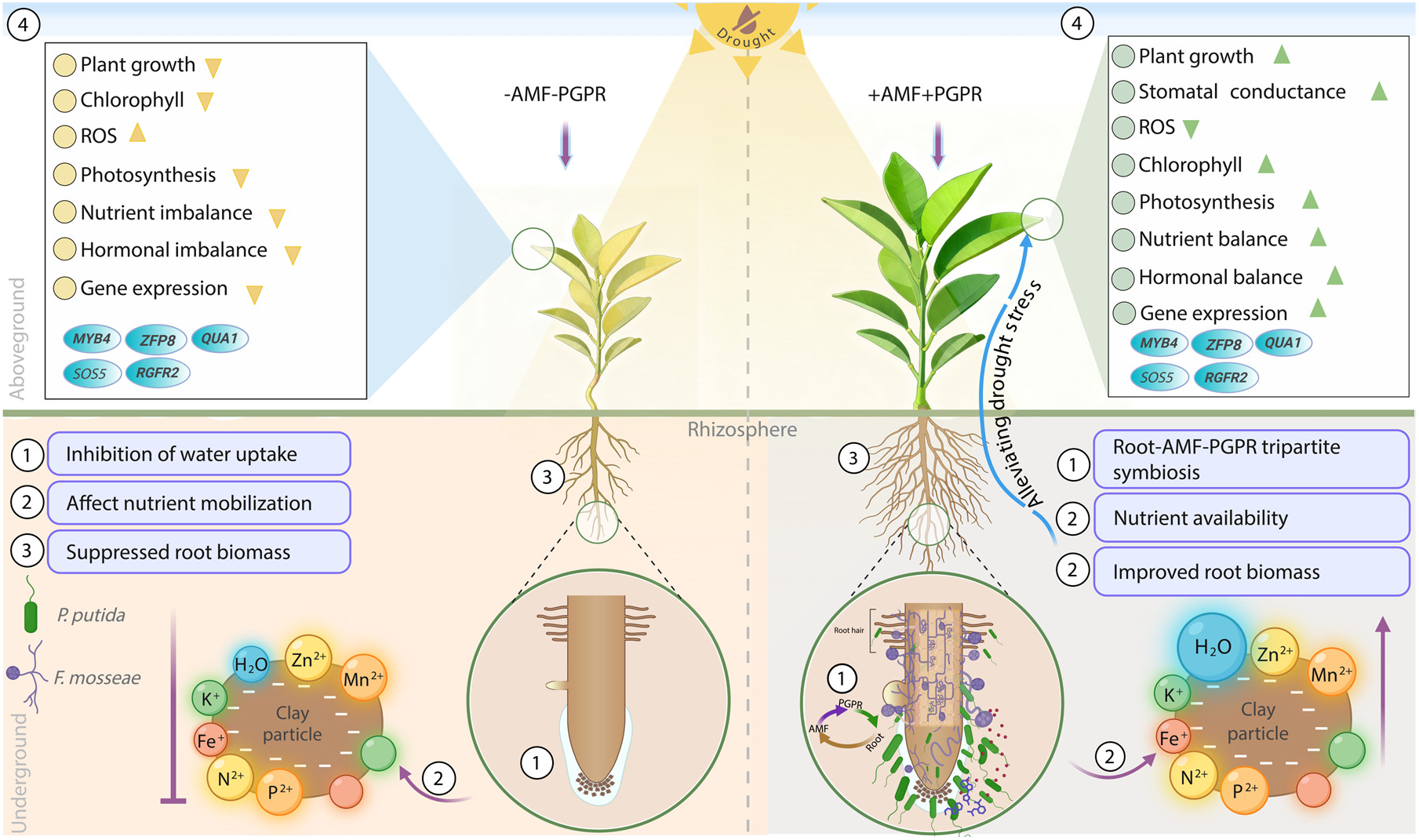
Conceptual model of the morphological, physiological, biochemical and molecular responses of *Citrus reticulata* to root-AMF-PGPR tripartite symbiosis under drought stress. **(1-3)** illustrate belowground responses, including root system development, water status and nutrient acquisition in inoculated versus non-inoculated plants. **(4)** Summarizes the corresponding aboveground responses (morphological, physiological, biochemical and molecular) in drought-stressed plants with or without microbial inoculation.

## Acknowledgements

The work was supported by the Chongqing Postdoctoral Research Project Special Funding of China (2022CQBSHTB1001), the National Foreign Expert Project of China (Y20240129), and the Postgraduate Research Innovation Project of Southwest University of China (SWUB23058).

## Author contributions

SU proposed the study and designed the experiments. SU and JY conducted and performed the experiments. SU and SG wrote the manuscript. SU analyzed RNA-Seq data. HAH, JW and UM revised the manuscript. XY supervised the study

## Data availability

The high-throughput sequencing data generated in this study, including RNA-seq datasets, have been deposited in the NCBI Sequence Read Archive (SRA) under accession number PRJNA1405962. Source data are available with this paper.

